# *ROSALIND* protects the mitochondrial translational machinery from oxidative damage

**DOI:** 10.1101/2024.01.04.574152

**Authors:** Vicky Katopodi, Yvessa Verheyden, Sonia Cinque, Elena Lara Jimenez, Ewout Demesmaeker, Alessandro Marino, Rita Derua, Elisabetta Groaz, Eleonora Leucci

**Affiliations:** Laboratory for RNA Cancer Biology, Department of Oncology, KU Leuven, Leuven, Belgium; Laboratory for Protein Phosphorylation and Proteomics; Rega Institute for Medical Research, Medicinal Chemistry, KU Leuven, Leuven, Belgium; Trace, Leuven Cancer Institute, KU Leuven, Leuven, Belgium

## Abstract

Upregulation of mitochondrial respiration coupled with high ROS-scavenging capacity is a characteristic shared by drug-tolerant cells in several cancers. As translational fidelity is essential for cell fitness, protection of the mitochondrial and cytosolic ribosomes from oxidative damage is pivotal. While mechanisms for recognizing and repairing such damage exist in the cytoplasm, the corresponding process in the mitochondria remains unclear.

By performing Ascorbate PEroXidase (APEX)-proximity ligation assay directed to the mitochondrial matrix followed by isolation and sequencing of RNA associated to mitochondrial proteins, we identified the nuclear-encoded lncRNA *ROSALIND* as an interacting partner of ribosomes. *ROSALIND* is upregulated in recurrent tumours and its expression can discriminate between responders and non-responders to immune checkpoint blockade in a melanoma cohort of patients. Featuring an unusually high G content, *ROSALIND* serves as a substrate for oxidation. Consequently, inhibiting *ROSALIND* leads to an increase in ROS and protein oxidation, resulting in severe mitochondrial respiration defects. This, in turn, impairs melanoma cell viability and increases immunogenicity both *in vitro* and *ex vivo* in preclinical humanized cancer models. These findings underscore the role of *ROSALIND* as a novel ROS buffering system, safeguarding mitochondrial translation from oxidative stress, and shed light on potential therapeutic strategies for overcoming cancer therapy resistance.

## Introduction

Despite all breakthroughs in targeted therapy and immune checkpoint blockade (ICB), cancer therapy resistance, whether pre-existing or arising following the acquisition of *de novo* mutations and via non-genetic adaptation, is still a clinical challenge. One of the most prominent non-genetic adaptation programs in the cell is the reprogramming of protein synthesis^1,2^, which is a prerequisite not only for tumour initiation and progression^3^ but also plays a crucial role in the generation of Drug Tolerant Persister Cells (DTPCs)^4,5^. In keeping with this, several intracellular and extracellular stressors converge to regulate the activity of cytosolic ribosomes to favour the translation of a specific subset of pro-survival/resistance mRNAs^1,3,6,7^. It was recently demonstrated that a large subset of the pro-survival mRNAs encodes for mitochondrial proteins, indicating that mitochondria are critical for the generation and survival of DTPCs^4^. In fact, as our understanding of DTPC biology increases, it is becoming evident that these cells exhibit a higher dependence on mitochondria than their drug-naïve counterparts, suggesting that mitochondrial biology is a very promising target for cancer therapy. As a matter of fact, antibiotics of the tetracycline family, which selectively block mitochondrial translation, prevented the emergence of most drug-tolerant subpopulations and delayed (and in some instances even abolished) the development of resistance to MAPK inhibitors in melanoma patient-derived xenograft (PDX) models^4^. Accordingly, resistance to multiple therapeutic modalities via the upregulation of mitochondrial respiration is a characteristic shared by several cancers^8,9^. A by-product of oxidative phosphorylation is Reactive Oxygen Species (ROS): a double-edged sword for the cell. At low levels ROS are essential signalling molecules, modifying phosphatases and metabolic enzymes and thus favour adaptation, however when increasing they induce peroxidation of proteins, lipids and nucleic acids that can irreparably compromise the functioning of the cell^10^. Oxidation of rRNA at the ribosomes’ catalytic centre was also shown to occur and to affect translation elongation^11^. In keeping with this, multiple anticancer drugs are thought to cause cell death in part via increasing the ROS cellular levels^12^. It follows that therapy resistant clones have robust systems to neutralise ROS^13–15^. Such systems are based on the conversion of superoxide to hydrogen peroxide (H_2_O_2_) by the enzymes of the SuperOxide Dismutases family (SOD). The resulting H_2_O_2_ is then converted to water by the CATalases (CAT), PeRoxireDoXins (PRDXs) and Glutathione PeroXidases (GPX) which undergo cycles of oxidation and reduction^10^.

Furthermore, the cell has also developed systems to recognise and replace damaged cellular components^16^. This is the case for oxidised cytosolic ribosomal proteins which are released from ribosomes through a chaperon-mediated ribosome repair mechanism^17^. Given the high levels of ROS in the mitochondria and the key role played by mitochondrial translation for cancer cell fitness^4,18,19^, it is unclear how oxidative damage is prevented at mitochondrial ribosomes. As it was demonstrated that lncRNAs play key role in the regulation of mitochondrial translation^18,19^ and more in general in melanoma mitochondrial biology^8^, we sought to investigate the dark genome further in search of lncRNAs interacting with the mitochondrial ribosome.

By performing Ascorbate PEroXidase (APEX)-proximity ligation assay directed to the mitochondrial matrix followed by isolation and sequencing of RNA associated with mitochondrial proteins, we identified LINC01918 as a mitoribosome-associated lncRNA protecting mitochondrial ribosomes from oxidative damage. This lncRNA, renamed *ROSALIND* for <u>RO</u>S <u>S</u>c<u>A</u>venging <u>Li</u>ncRNA mitocho<u>nd</u>rial, represents a previously unrecognized ROS scavenging system and a potential therapeutic target.

## Results

### *ROSALIND* is a nuclear-encoded lncRNA localising in the mitochondrial matrix

To identify lncRNAs implicated in the regulation of mitochondrial translation we performed APEX proximity biotinylation of mitochondrial matrix proteins following block mitochondrial protein synthesis (Figure 1A) by sublethal (Supplementary figure 1A-B) Tigecycline treatment. RNAs bound to the biotinylated proteins were then identified by streptavidin purification and bulk RNA-Sequencing. The downregulation of mitochondrial translation and the efficiency of the pull-down were confirmed by western blot (WB) for streptavidin (Figure 1B). The purity of the pull-down was confirmed by WB for mitochondrial (ATP5A, NDUSF3, TOM20), nuclear (XRN2) and cytoplasmic (vinculin) targets (Figure 1B). The sequencing yielded 920 significantly depleted transcripts upon tigecycline treatment and 1030 significantly enriched transcripts (Figure 1C), 10.6% to 13.6% of them being ncRNAs. Among the lincRNAs regulated upon the block of mitochondrial translation, we identified 2 transcripts significantly enriched in the mitochondrial matrix and 7 depleted (Figure 1D), including *SAMMSON* (p=0.02 log2FC=-1.46)^18,19^ and *LENOX* (p=0.009 log2FC -1.40)^8^, previously reported to localise at mitochondria. Out of them, LINC01918 (AC104655.3), from now on *ROSALIND,* attracted our attention as it was the second most depleted lincRNA upon tigecycline (adj p=7.17e-19 log2FC=-3.21) (Table 1 and Figure 1D). *ROSALIND* gene is located on Chromosome 2, ∼44kb bases downstream of the locus encoding for the mitochondrial ribosomal protein MRPS9 and amplified in 63% of melanoma cases and coamplified with MRPS9 in 99% of cases (Figure 1E-F). Analysis by qPCR of the different *ROSALIND* isoforms retrieved in the APEX experiment (Supplementary figure 1C) suggested that ENST00000436841 is the polyadenylated isoform enriched at mitochondria (Supplementary figure 1D-E). *ROSALIND* localisation to the mitochondria was also confirmed by cell fractionation (Figure 1G), where *SAMMSON* and the mitochondrial rRNA 16S were used as a positive control, and by Fluorescent In Situ Hybridization (FISH) colocalisation with the mitochondrial protein p32 (Figure 1H-I). In keeping with the APEX results, decreased expression of *ROSALIND* upon tigecycline treatment was detected by quantifying the FISH images (Figure 1J) and by qPCR in 3 different melanoma cell lines (Supplementary figure 1F), while no significant change in the nuclear/cytoplasmic distribution of *ROSALIND* occurred upon tigecycline treatment (Figure 1K). These data demonstrated that *ROSALIND* is a lncRNA residing in the mitochondrial matrix and possibly linked to mitochondrial translation.

**Figure 1A-K:**
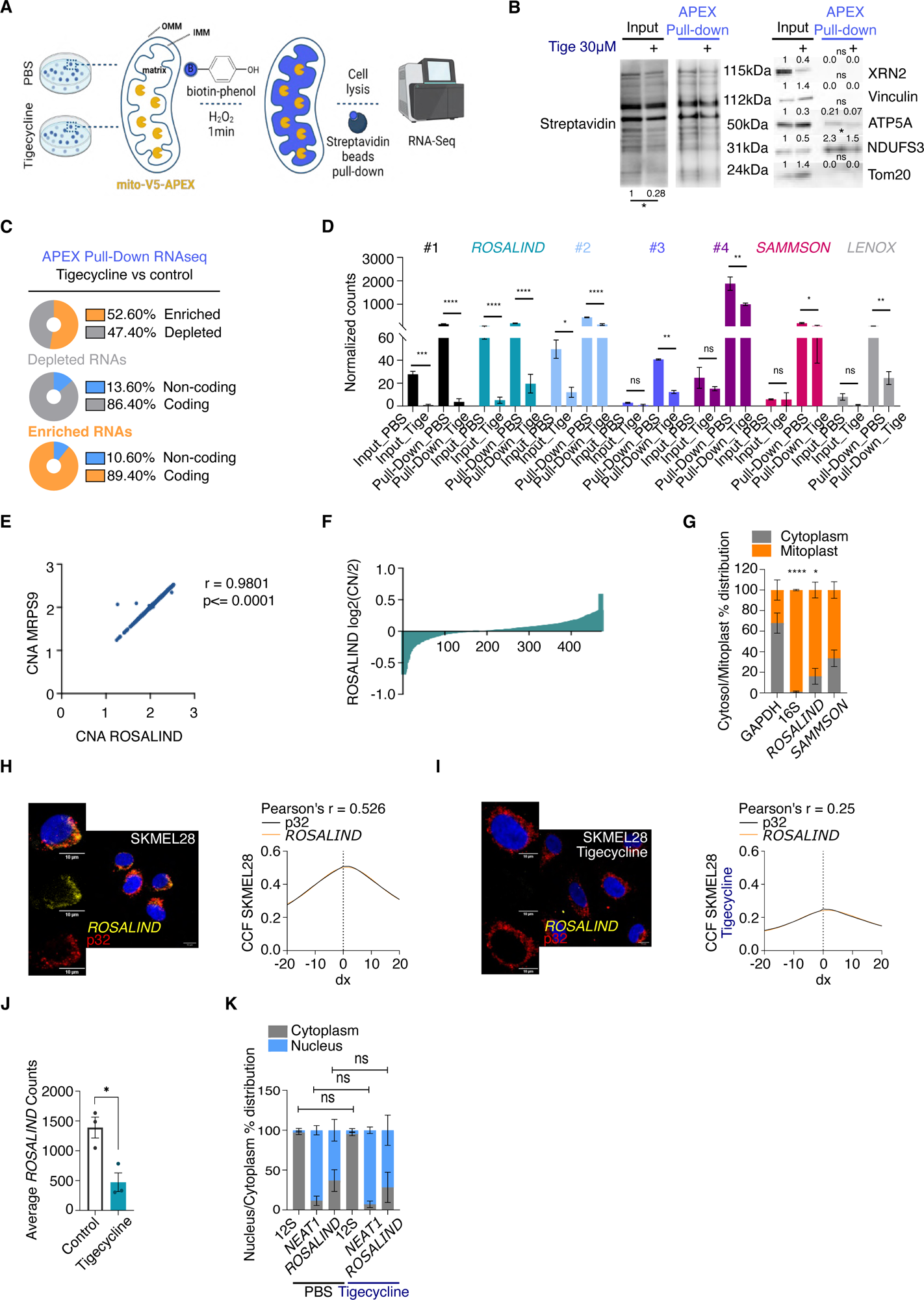
*ROSALIND* is a nuclear-encoded lncRNA localising to the mitochondrial matrix. **A.** Schematic representation of APEX-Seq workflow performed on SKMEL28 cells **B.** Evaluation of the efficiency of Biotin pulldown by WB for streptavidin (left) Evaluation of the APEX pulldown efficiency and specificity (right) by WB for different cellular markers. **C.** Analysis of the RNA retrieved in the matrix upon block of mitochondrial translation as assessed by APEX-Seq. **D.** Counts of lncRNAs retrieved in the APEX-Seq input and pull-down normalised on library size as calculated by DESeq2 differential expression analysis. **E.** Correlation between MRPS9 and *ROSALIND* CNA in the TCGA melanoma cohort. Significance has been calculated by Spearman rank correlation analysis **F.** DNA copy number of *ROSALIND* and MRPS9 in melanoma lesions from the TCGA melanoma cohort. Samples are ranked according to ROSALIND copy number and expressed as the mean log ratio. **G.** *ROSALIND* cytosolic/mitochondrial ratios (in percentage) as assessed by RT-qPCR on mitoplast extracts of SKMEL28 cells. *SAMMSON* and the mitochondrial rRNA *16s* were used as positive controls. Paired t-test was used to calculate statistical significance between cytosol and mitoplast localisation. **H.** Left panel: representative confocal images from SKMEL28 control cells. *ROSALIND* is in yellow and nuclei are stained with DAPI (blue). P32 (in red) was used as a mitochondrial marker. Right panel: Evaluation of colocalization between *ROSALIND* and p32 was measured by Cross Correlation Function (CCF) with a pixel shift of δ = ±20. ρ indicates Pearson’s coefficient. **I.** Representative confocal images from SKMEL28 cells treated with 30μM tigecycline for 24h. *ROSALIND* is in yellow and nuclei are stained with DAPI (blue). P32 (in red) was used as a mitochondrial marker. Evaluation of colocalization between *ROSALIND* and p32 was measured by Cross Correlation Function (CCF) with a pixel shift of δ = ±20. ρ indicates Pearson’s coefficient. **J.** Average *ROSALIND* counts in the images in H. and I. Significance was calculated by paired t-test. **K.** *ROSALIND* nuclear/cytoplasmic ratios (in percentage) in Ctrl and Tigecycline treated SKMEL28 as assessed by RT-qPCR. Significance was calculated by paired t-test.

**Table 1:**
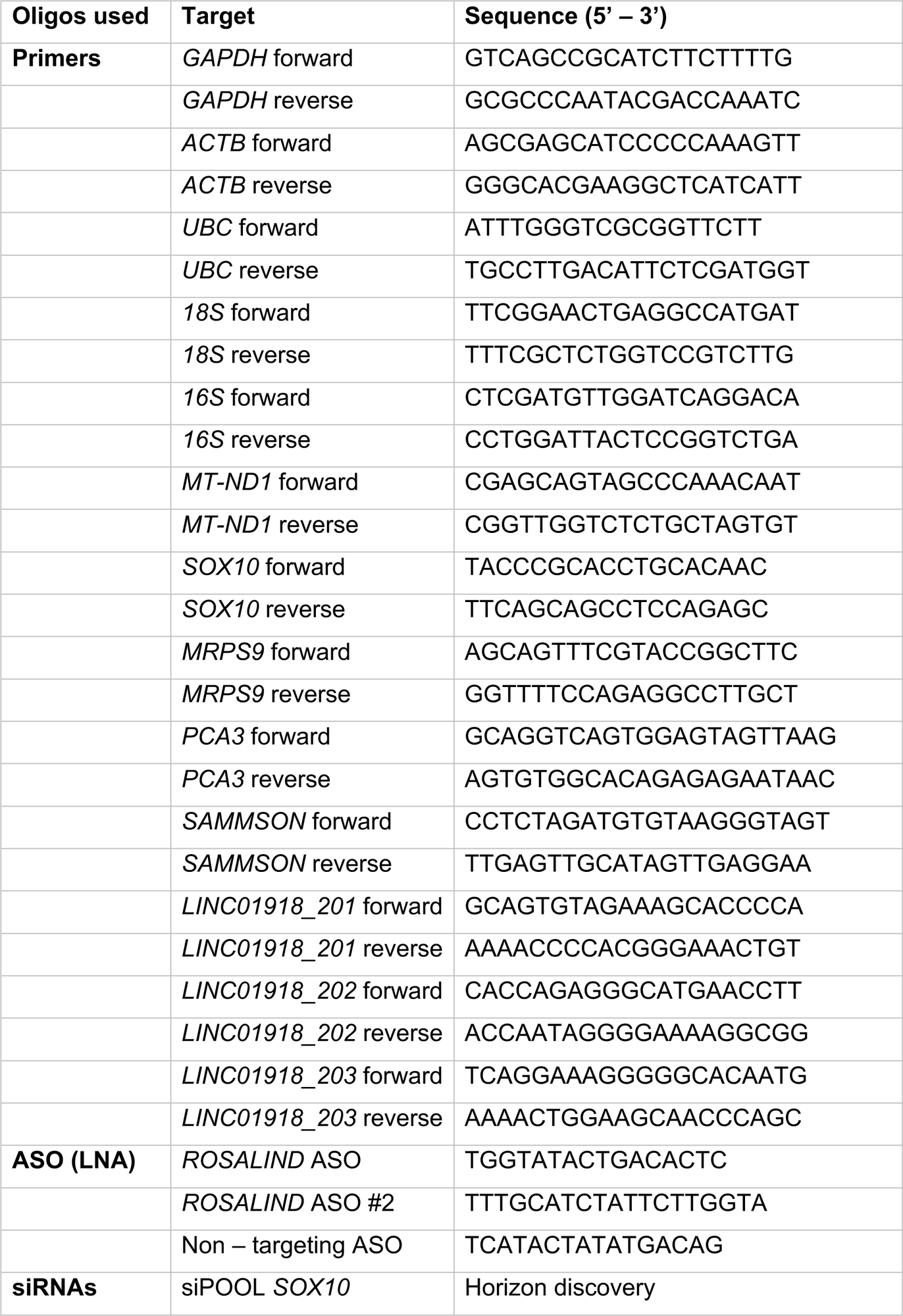
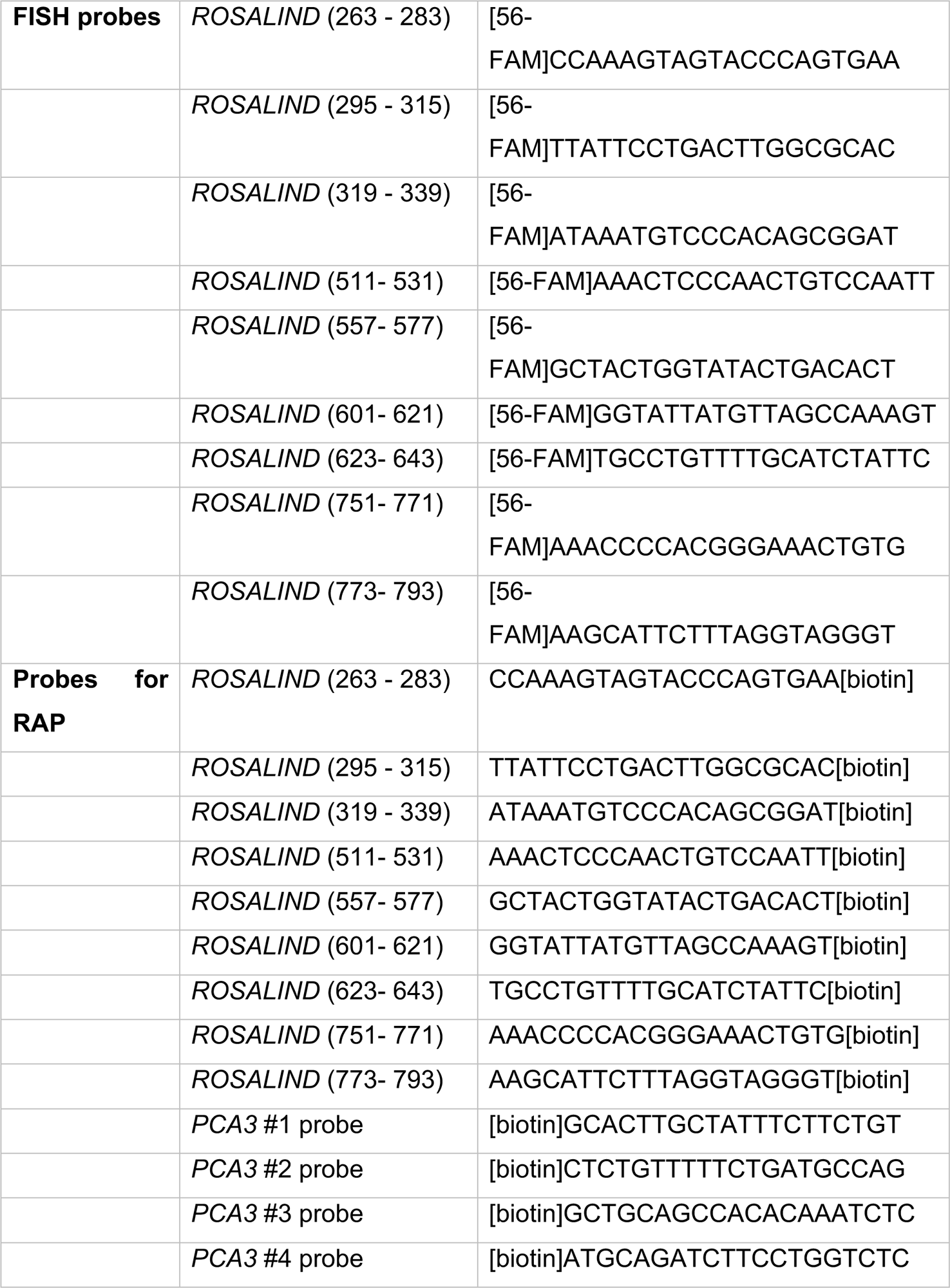
Sequences of primers, ASOs and probes.

### *ROSALIND is* expressed in normal skin and associated with recurrence and therapy resistance in melanoma

To better characterise *ROSALIND,* we analysed its expression in normal tissues from the Genotype-Tissue Expression (GTEx) database and in the PanCancer atlas. In normal conditions, *ROSALIND* is expressed in different areas of the brain and in the skin (Figure 2A). Analysis of the PanCancer atlas suggests however, that *ROSALIND* is broadly expressed in cancer (Figure 2B) where its expression was significantly higher in recurrent tumours (Figure 2C) and correlated with survival (Supplemental figure 1G). In melanoma, its expression could be detected in several melanoma lines irrespectively of their driver mutation and phenotypic state (Supplemental figure 1H). In keeping with this, *ROSALIND* was also expressed in a panel of PDX melanoma models derived from drug-naïve or drug-resistant patients (Supplemental figure 1I). Furthermore, a significantly higher expression of *ROSALIND* was detected in PDX models upon treatment with targeted therapy at relapse and in the same PDX model (resistant to immune checkpoint blockade) upon treatment with anti-PD1 (Figure 2D). These data were also further confirmed by FISH in the same PDX models (Figure 1E-H) where co-localisation with mitochondrial markers could also be detected (Supplementary Figure 1J-K). Similar results were also obtained in a cohort of patients from the SPECIAL clinical trial from UZ Leuven^20^ where scRNAseq (scRNAseq) was performed before and early on treatment with immunotherapy (Figure 2I). Here we could detect *ROSALIND* expression selectively in patients non-responding to the therapy (Figure 2I) in which no immune infiltration could be detected (Figure 2J). In keeping with this, guilty by association analysis by Gene Set Enrichment Ananlysis (GSEA) suggested a role of *ROSALIND* in OXPHOS and a negative correlation with immune responses (p value<0.05 and FDR<0.1; Figure 2K). These data demonstrated that *ROSALIND* expression correlates with patient survival and therapy responses.

**Figure 2A-L:**
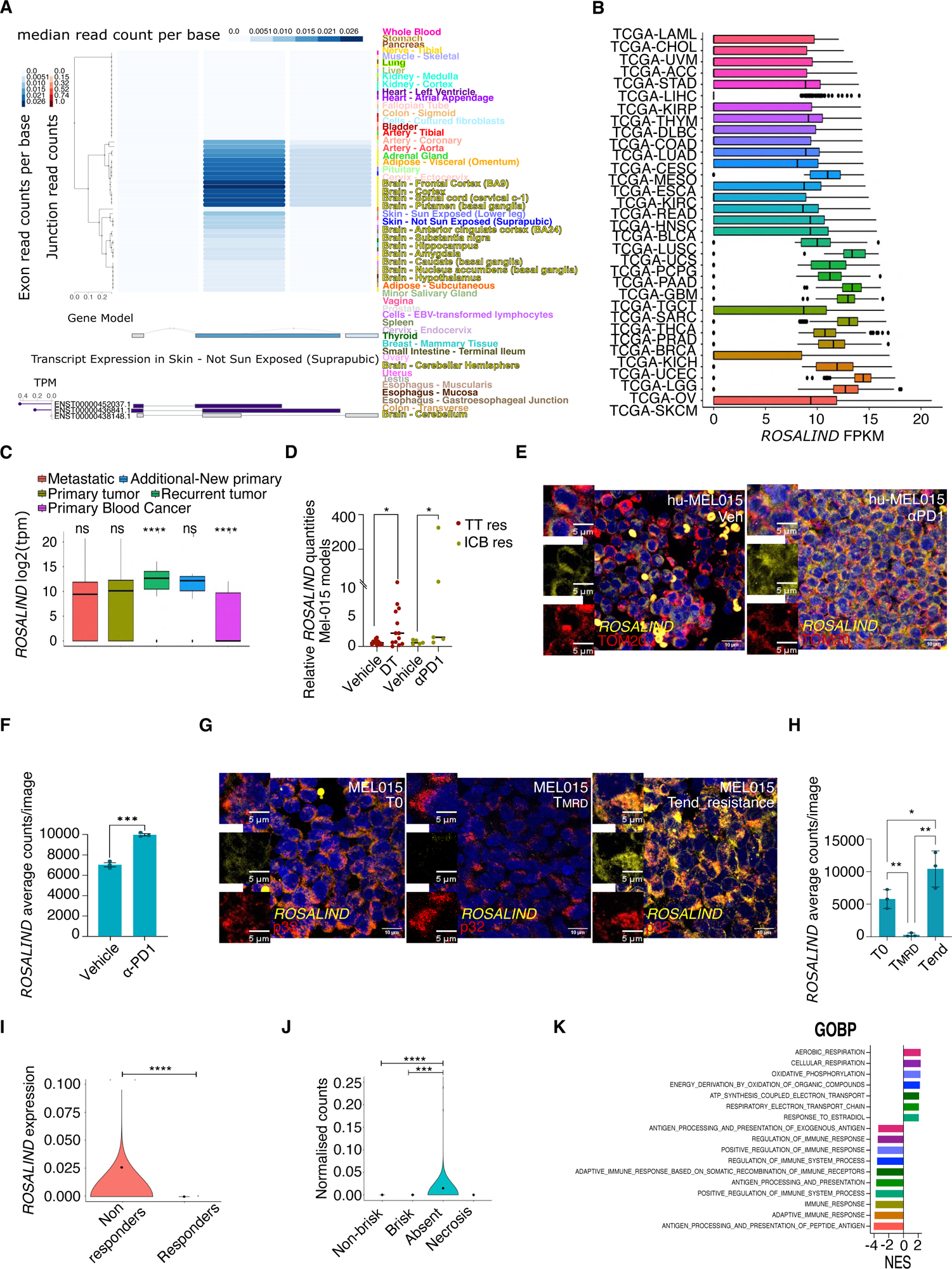
*ROSALIND* expression correlates with cancer recurrence and therapy resistance. **A.** *ROSALIND* expression in a panel of adult normal samples from the GTEx database. **B.** Expression of *ROSALIND* in the PanCancer Atlas across different tumours. **C.** Expression of *ROSALIND* in the PanCancer Atlas per type. **D.** Relative *ROSALIND* expression as assessed by qPCR in Mel-015 PDX at relapse from targeted therapy and after treatment with α-PD1 (non-responder). Significance was calculated by unpaired t-test. **E.** Representative confocal images from huMEL015 PDX models treated with immunotherapy (α-PD1) or control substance, at the end of treatment. *ROSALIND* is in yellow and nuclei are stained with DAPI (blue). TOM20 (in red) was used as a mitochondrial marker. **F.** Average *ROSALIND* counts from the experiment shown in E. calculated using ImageJ. Each dot represents the average counts of each PDX used. Paired t-test was used to calculate significance. **G.** Representative confocal images from MEL015 PDX models at the start (T0), MRD phase (T_mrd) and relapse (Tend_resistance) of targeted therapy (Dabrafenib-Trametinib). *ROSALIND* is in yellow and nuclei are stained with DAPI (blue). P32 (in red) was used as a mitochondrial marker. **H.** Average *ROSALIND* counts from the experiment shown in G. calculated using ImageJ. Each dot represents the average counts of each PDX used. Paired t-test was used to calculate significance **I.** *ROSALIND* expression (read counts normalised by Sleuth) in a melanoma cohort of responders and not responders to immune checkpoint blockade^20^. Significance was calculated by Mann Whitney test. **J.** Correlation of *ROSALIND* expression with tumour infiltration status in patients from clinical trial described in I. Significance was calculated by Kruskal-Wallis test. **K**. GSEA of the transcript co-expressed with *ROSALIND* in the patient cohort in I.

### *ROSALIND* binds to the mitochondrial ribosome and affects respiration

To find out whether *ROSALIND* plays a role in the regulation of mitochondrial translation, we performed puromycin-labelling of cytosolic and mitoplast newly synthetized proteins, followed by mitoplast (mitochondria stripped of their outer membrane) fractionation. Western blot analysis of puromycin incorporation in Mel-015 melanoma cells that acquired resistance to targeted therapy, revealed that depletion of *ROSALIND* (Figure 3A) inhibits mitochondrial translation, thus suggesting a role in protein synthesis inside the organelle (Figure 3B-C). To check whether this was mediated by regulation of MRPS9 *in cis,* since *ROSALIND* is also detectable in the nucleus of melanoma cells, we silenced it in four different melanoma lines and performed qPCR for MRPS9, which was found to be unaffected (Supplemental figure 2A). Therefore, to investigate the function of *ROSALIND* in an unbiased way, we thought to identify its protein partners in tumours derived from a PDX models of acquired resistance to targeted therapy (DT: Dabrafenib plus Trametinib), as these models are well known to have high OXPHOS and increased mitochondrial dependency at resistance. We performed RNA Affinity Purification (RAP) coupled to Mass spectrometry in vehicle- and DT-treated tumours (Supplemental Table 1). Analysis of the protein content of these tumours by GSEA, detected enrichment in OXPHOS activity and reduction in cytosolic translation rates in resistant tumours, compatibly with their dependence on mitochondria (Supplemental figure 2B). Among the protein interactors retrieved both in control conditions and upon therapy-resistance acquisition, we found the mitochondrial ribosomal protein MRPL38 (Figure 3D). MRPL38 is part of the large ribosomal subunit, and it is positioned at the top of the ribosomal central protuberance, where peptide bond formation occurs^21^. The interaction was also validated by RAP-western blot (Figure 3E) and by RNA Immune precipitation (RIP) using single cell suspension generated from tumour extracts at T0 or after targeted therapy resistance acquisition (Figure 3F-H). To investigate whether *ROSALIND* associates with the mitochondrial ribosome or with ribosome-free MRPL38, we performed ribosome profiling on mitoplast extracts from melanoma cells and confirmed that *ROSALIND* was significantly enriched in the large mitochondrial ribosomal subunit (39S) and the mitochondrial monosome (Figure 3I-J). To rule out the possibility that *ROSALIND* associates with mitochondrial ribosomes to be translated, we checked its predicted coding potential using the Coding Potential Calculator (CPC2) algorithm (Supplemental figure 2C). Additionally, we could not find peptides derived from this transcript in mass spec data generated by isolated mitoplast extracts of melanoma cells^7^. Knowing that silencing of *ROSALIND* reduced mitochondrial translation, which is responsible for the generation of proteins that take part in the assembly of electron transport chain complexes (ETC), we measured oxygen consumption rates in a cell line derived from the above-mentioned Mel-015 model that acquired resistance to targeted therapy, upon *ROSALIND* KD. *ROSALIND* depletion (Figure 3K) caused a marked and significant decrease in both basal and maximal respiration as well as in ATP production (Figure 3L-M). Accordingly, imaging of mitochondria stained with MitoTracker Red and subsequent measurement of different parameters (shape, number, mass…), in melanoma cells resistant to targeted therapy upon *ROSALIND* KD (Figure 3N) revealed changes in mitochondrial morphology and decrease in their number, supportive of severe mitochondrial dysfunction (Figure 3O-P). Conversely, no obvious changes in mitochondria branching and lenght could be detected (Supplemental figure 2D). These data could also be validated with one additional ASO (Supplemental figure 2E-G). Similar changes, albeit less profound, were also observed in drug naïve melanoma cells of the melanocytic lineage (Supplemental figure 2H-J). To better understand the contribution of *ROSALIND* to mitochondrial translation, we overexpressed it in SKMEL28 melanoma cells, and checked translation rates by puromycin incorporation assay (Figure 3Q-S). However, despite the correct *ROSALIND* localization to the mitochondrial matrix (Supplemental figure 2K), the overexpression (Figure 3Q) failed to produce any change in the translation rates in the mitochondria or in the cytosol, indicating that *ROSALIND KD* effects on translation are the results of ETC defects (Figure 3R-S). Collectively, these data demonstrate that *ROSALIND* binds to the mitoribosomes and affects mitochondrial functions.

**Figure 3A-S:**
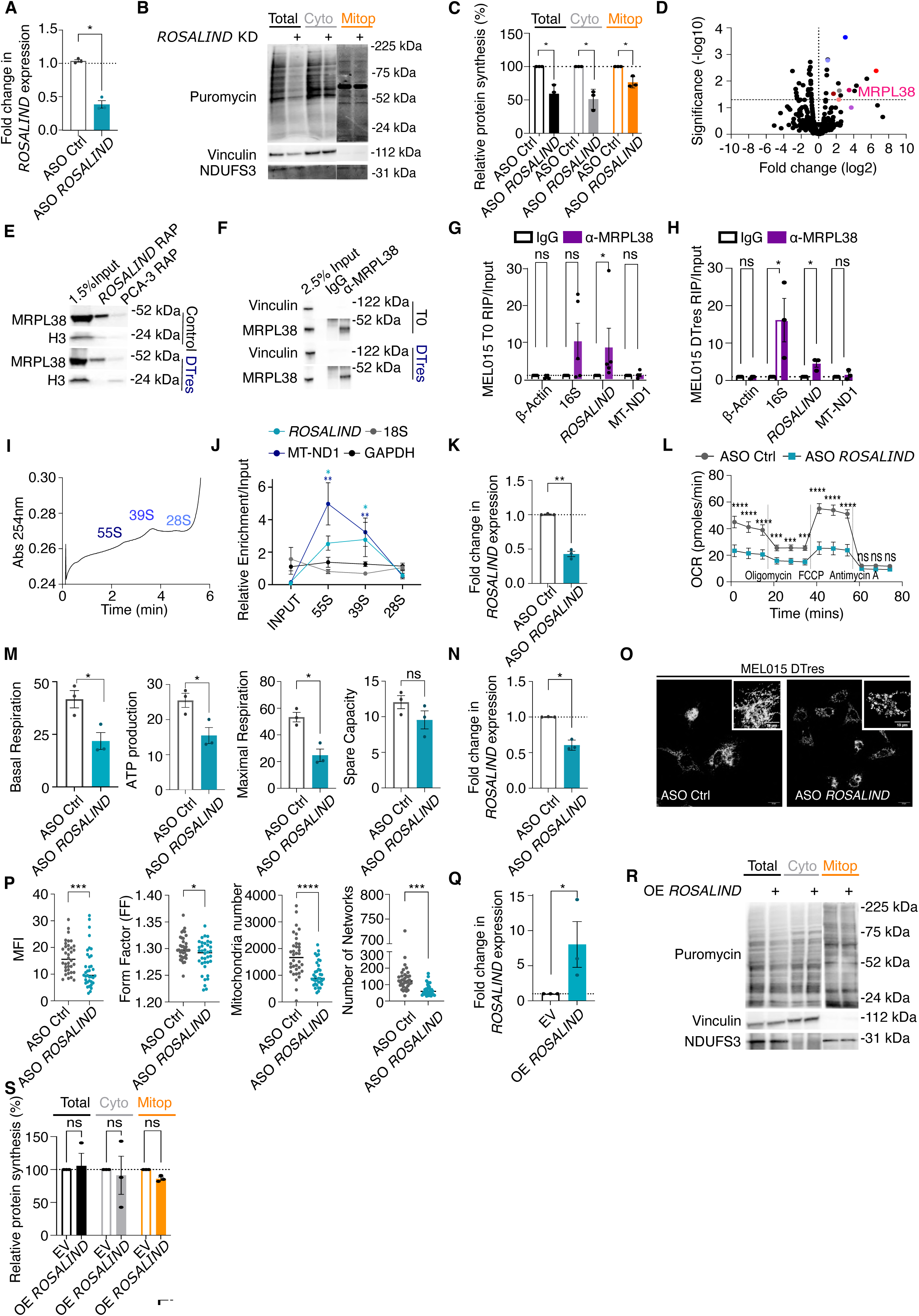
*ROSALIND* affects mitochondrial mass and respiration. **A.** Fold changes in *ROSALIND* expression upon its KD in the PDX derived melanoma cell line MEL015 upon acquisition of therapy resistance (MEL015_DTres) as calculate by qPCR. Significance was calculated by paired t-test. **B.** WB assessment of a puromycin incorporation assay performed on whole cell lysates, cytoplasmic and mitoplast extracts upon *ROSALIND* KD in MEL015_DTres depicted in A. **C.** Densitometry of the western blot in B. Paired t-test was used to calculate statistical significance. **D.** Volcano plot illustrating *ROSALIND* protein partners identified by RAP-Mass Spectrometry in DT-treated PDX tumours at relapse. A probe set against *PCA-3* was used as a negative control. The log_2_ fold change is plotted against statistical significance. **E.** Validation of *ROSALIND* interaction with MRPL38 by RAP-WB. H3 is used as loading control. **F.** Efficiency of MRPL38 RIP as assessed by WB. Vinculin was used as a loading control. **G.H.** *ROSALIND* enrichment over input in MRPL38 RIP samples derived from tumours isolated from vehicle-treated (G.) and DT treated (H.) MEL015 PDX model, as assessed by qPCR. 16S was used as a positive control. Significance was calculated by multiple paired t-test. **I.** Absorbance of Mitoribosome fractions obtained upon ultracentrifugation of mitoplast extracts loaded on sucrose gradients. The position of the different subunits and of the mitochondrial monosome (55S) as calculated from rRNA enrichments is indicated in the graph. **J.** Expression of *ROSALIND* in mitoribosome fractions as assessed by RT-qPCR. The mitochondrial protein coding RNA MT-ND1 is shown as a positive control. 18S and GAPDH were used as negative controls. Paired t-test was used to calculate statistical significance. **K.** Fold changes in *ROSALIND* expression upon its KD in MEL015_DTres as calculated by qPCR. Significance was calculated by paired t-test. **L.** Oxygen consumption rates of MEL015_DTres cells upon inhibition of *ROSALIND* as measured by Seahorse. Significance was calculated by two-way ANOVA mixed effect. **M.** Basal respiration, ATP production, maximal respiration and spare capacity were calculated as an average of three values taken by the Seahorse before the addition of each inhibitor. Each dot represents a biological replica. Significance was calculated by Paired t-test. **N.** Fold changes in *ROSALIND* expression upon its KD in MEL015_DTres as calculate by qPCR. Significance was calculated by paired t-test. **O.** Representative confocal images from Mitotracker Red stained MEL015_DTres cells. **P.** Quantification of mitochondrial properties from the images shown in O. Calculation were performed with MiNa workflow on ImageJ. Each dot represents an image acquired from the confocal with multiple cells, that have been averaged for each of the mitochondrial properties. Images used for these graphs are derived from a biological triplicate. Significance was calculated by unpaired t-test. **Q.** Fold changes in *ROSALIND* expression upon its overexpression (OE) in SKMEL28 as calculate by qPCR. Significance was calculated by paired t-test. **R.** WB assessment of a puromycin incorporation assay performed on whole cell lysates, cytoplasmic and mitoplast extracts upon *ROSALIND* KD in SKMEL28 depicted in Q. **S**. Densitometric analysis of the WB in R. Paired t-test was used to calculate statistical significance.

### *ROSALIND* depletion has cytotoxic effects caused by aberrant ROS production

Since mitochondria are important in DTPC biology, we sought to investigate the role of *ROSALIND* on cellular fitness *in vitro*, using a panel of established melanoma cell lines that phenotypically resemble these DTPCs (Supplemental figure 1H). We selected therefore four cell lines with different driver mutations and different *ROSALIND* expression levels and transfected them with a non-targeting control ASO (Ctrl ASO) or an ASO targeting *ROSALIND. ROSALIND* knock-down (Figure 4A) led decreased cell confluency in both MM383 and MM047 (Figure 4B, D, E, G) and lead to a significant increase in cell death irrespective of driver mutation (Figure 4B, C, E, F). Importantly, no change in confluency (Supplemental figure 3A, B) or cell death (Supplemental figure 3A,C) were detected in HeLa cells that do not express *ROSALIND* (Supplemental figure 3D), thus ruling out possible ASO toxicity. Surprisingly however, *ROSALIND* KD had no deadly effect on MM099 (Figure 4H-J) and SK-MEL-28 (Figure 4K-M) cell lines, which showed only reduced growth rates. When comparing the gene expression profiles of responders and non-responders^4,22^ by GSEA, we detected an enrichment in oxidoreductase activity in the cell lines not responding to the *ROSALIND* depletion (Figure 4N). This was further supported by the higher catalase levels detected by WB in these cells (Supplementary figure 3E-F). Based on the above, we hypothesised that the increase in ROS was the cause of melanoma death and that MM099 and SKMEL28 cell lines possibly had a more robust ROS buffering capacity, preventing cell death upon *ROSALIND* deprivation. In keeping with this, flow cytometry of melanoma cell lines subjected to *ROSALIND* depletion (Supplemental figure 3G) and stained with MitoSOX Red, a fluorescent dye used to measure mitochondrial ROS, revealed a significant increase in their levels in the absence of *ROSALIND* in all the three cell lines tested (Figure 4O, Supplemental figure 3H-J). Furthermore, the addition of the antioxidant N-acetyl-L-cystein (NAC) rescued the cell viability phenotype in melanoma cell lines sensitive to *ROSALIND* depletion but had no effect on MM099 (Figure 4O-P and Supplemental figure 3H-J). These data were further validated with an independent ASO (Supplemental figure 3K-Q). Conversely, overexpression of *ROSALIND* in MM383 and in MM099 (Supplemental figure 4A) -respectively affected and not affected by *ROSALIND* KD-successfully decreased ROS release (Supplemental figure 4B-D) and improved cell viability selectively in MM383 cells (Supplemental figure 4F-G). Lastly, overexpression of *ROSALIND* further increased resistance to targeted therapy in MEL015_DTres line. Overall, these data suggest that *ROSALIND* act as a novel ROS scavenging system.

**Figure 4A-R:**
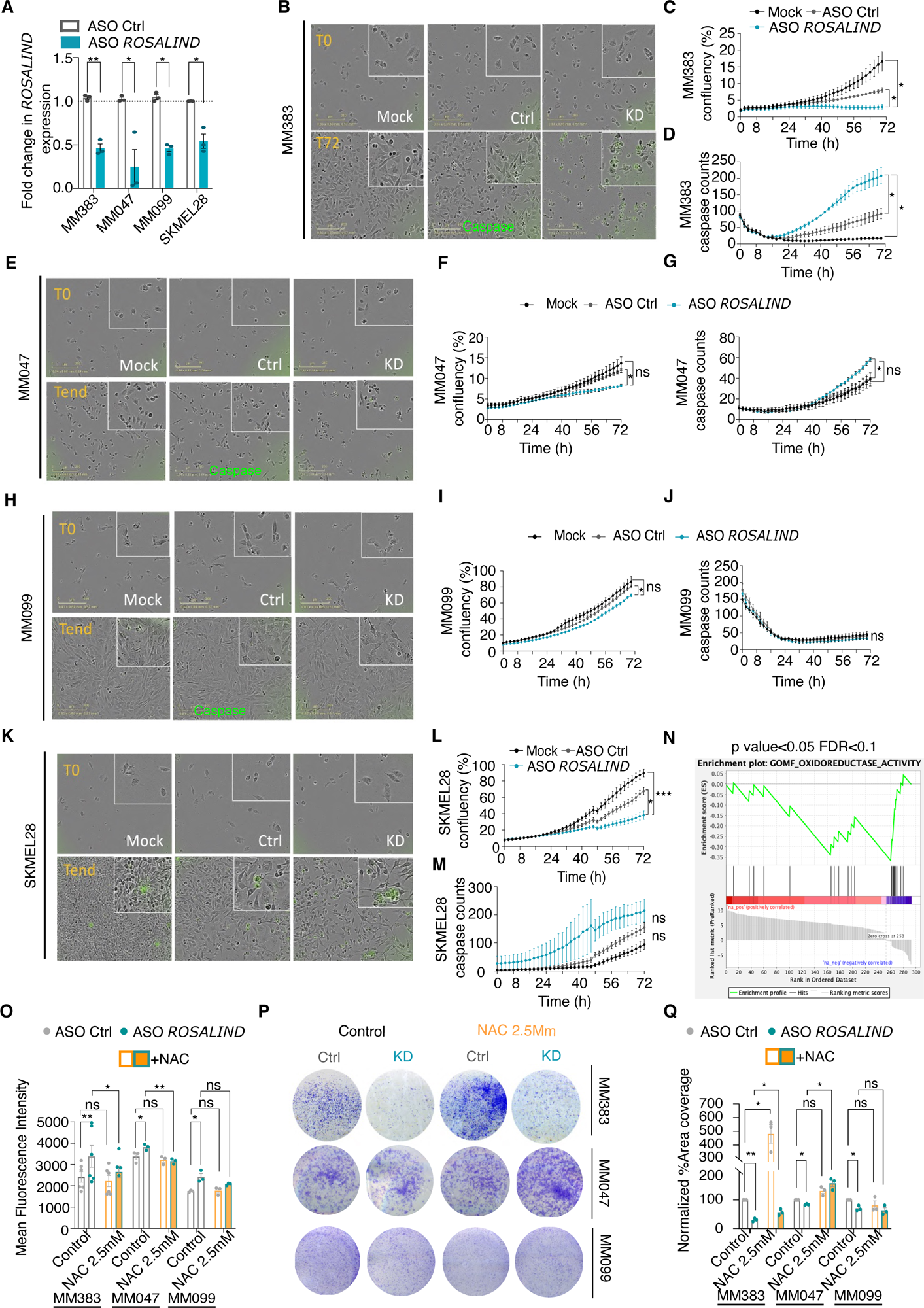
*ROSALIND* affects melanoma viability. **A**. Fold changes in *ROSALIND* expression upon its KD in different melanoma lines. Significance was calculated by paired t-test. **B.** Representative Incucyte pictures from MM383 cells transfected with ASO *ROSALIND*, ASO control (ASO Ctrl) or not transfected (Mock) at T_0_ and T_72_ hours. Caspase-3+ cells are in green **C.** Analysis of MM383 confluency (%) transfected as indicated in B. Two-way ANOVA mixed effect was used to calculate statistics **D.** Analysis of Caspase-3 counts in MM383 transfected as indicated in B. Two-way ANOVA mixed effect was used to calculate statistics. **E.** Representative Incucyte pictures from MM047 cells transfected with ASO *ROSALIND*, ASO control (ASO Ctrl) or not transfected (Mock) at T_0_ and T_72_ hours. Caspase-3+ cells are in green **F.** Analysis of MM047 confluency (%) transfected as indicated in E. Two-way ANOVA mixed effect was used to calculate statistics. **G.** Analysis of Caspase-3 counts in MM047 transfected as indicated in E. Two-way ANOVA mixed effect was used to calculate statistics. **H.** Representative Incucyte pictures from MM099 cells transfected with ASO *ROSALIND*, ASO control (ASO Ctrl) or not transfected (Mock) at T_0_ and T_72_ hours. Caspase-3+ cells are in green **I.** Analysis of MM099 confluency (%) transfected as indicated in H. Two-way ANOVA mixed effect was used to calculate statistics **J.** Analysis of Caspase-3 counts in MM099 transfected as indicated in H. Two-way ANOVA mixed effect was used to calculate statistics. **K.** Representative Incucyte pictures from SKMEL28 cells transfected with ASO *ROSALIND*, ASO control (ASO Ctrl) or not transfected (Mock) at T_0_ and T_72_ hours. Caspase-3+ cells are in green. **L.** Analysis of SKMEL28 confluency (%) transfected as indicated in K. Two-way ANOVA mixed effect was used to calculate statistics **M.** Analysis of Caspase-3 counts in SKMEL28 transfected as indicated in K. Two-way ANOVA mixed effect was used to calculate statistics. **N.** GSEA on differentially expressed genes in cell lines responding and not responding to *ROSALIND* KD. Threshold values FDR<0.1 p<0.05. **O.** Evaluation of ROS production by MM383, MM047 and MM099 upon *ROSALIND* KD with (orange bars) and without (grey bars) NAC buffering, as measured by MitoSOX staining (mean fluorescent intensity) and flow cytometry analysis. Significance was calculated by two-way ANOVA. **P.** Colony assays of MM383, MM047 and MM099 upon *ROSALIND* KD with and without NAC buffering. **Q.** Quantification of colony assays in P expressed as percentage of area covered normalised to the Control sample (treated only with ASO Ctrl, without NAC). Two-way ANOVA mixed effect was used to calculate statistics.

### *ROSALIND* is a new ROS scavenging system

We next analysed carefully *ROSALIND*’s sequence and compared it to the sequence of 6123 ribosome-bound lncRNAs identified in patient-derived samples, to the identified in the mitochondria matrix by APEX, to mitochondrial protein-coding transcripts and to a general random set of lncRNAs. Our analysis revealed that *ROSALIND* had an overall significantly higher G content compared to all the subsets tested (Figure 5A). As G nucleotides are the ones that are predominantly modified by ROS, this data indicated that mitochondrial RNAs are intrinsically protected from oxidative damage by their low G content and that *ROSALIND* may act as a ROS scavenger to safeguard the formation of the peptide bond and the ribosomal proteins instead. For our hypothesis to be correct, we reasoned that *ROSALIND* should be heavily modified itself by ROS and furthermore its depletion should lead to an increase in oxidative damage in the mitochondrial matrix. To explore this possibility, we immunoprecipitated oxidised Gs from total RNA isolated from three different melanoma cell lines using the 8OHG antibody (Supplemental figure 5A) and measured the *ROSALIND* enrichment by qPCR. *ROSALIND* but not *LENOX* (which has low G content) was found consistently enriched in the IPs, indicating that it is oxidised in all the cell lines tested (Figure 5B). Furthermore, western blotting for protein carbonylation -a common product of oxidative damage-upon *ROSALIND* depletion (Figure 5C) followed by mitoplast isolation, confirmed a significant increase in protein damage only in the cell lines that show apoptotic death upon its KD (Figure 5D-E). These findings could be reproduced also with one additional ASO (Supplemental figure 5B-D). Knowing the association of *ROSALIND* with MRPL38, we performed knock down of *ROSALIND* in the PDX-derived cell line Mel-015 (Figure 5F) followed by immunoprecipitation of MRPL38 (Figure 5G-H) and western blot for carbonyl groups detected significantly more damage in MRPL38 (Figure 5I-J) suggesting that indeed *ROSALIND* is a novel ROS scavenging system in the mitochondrial matrix.

**Figure 5A-J:**
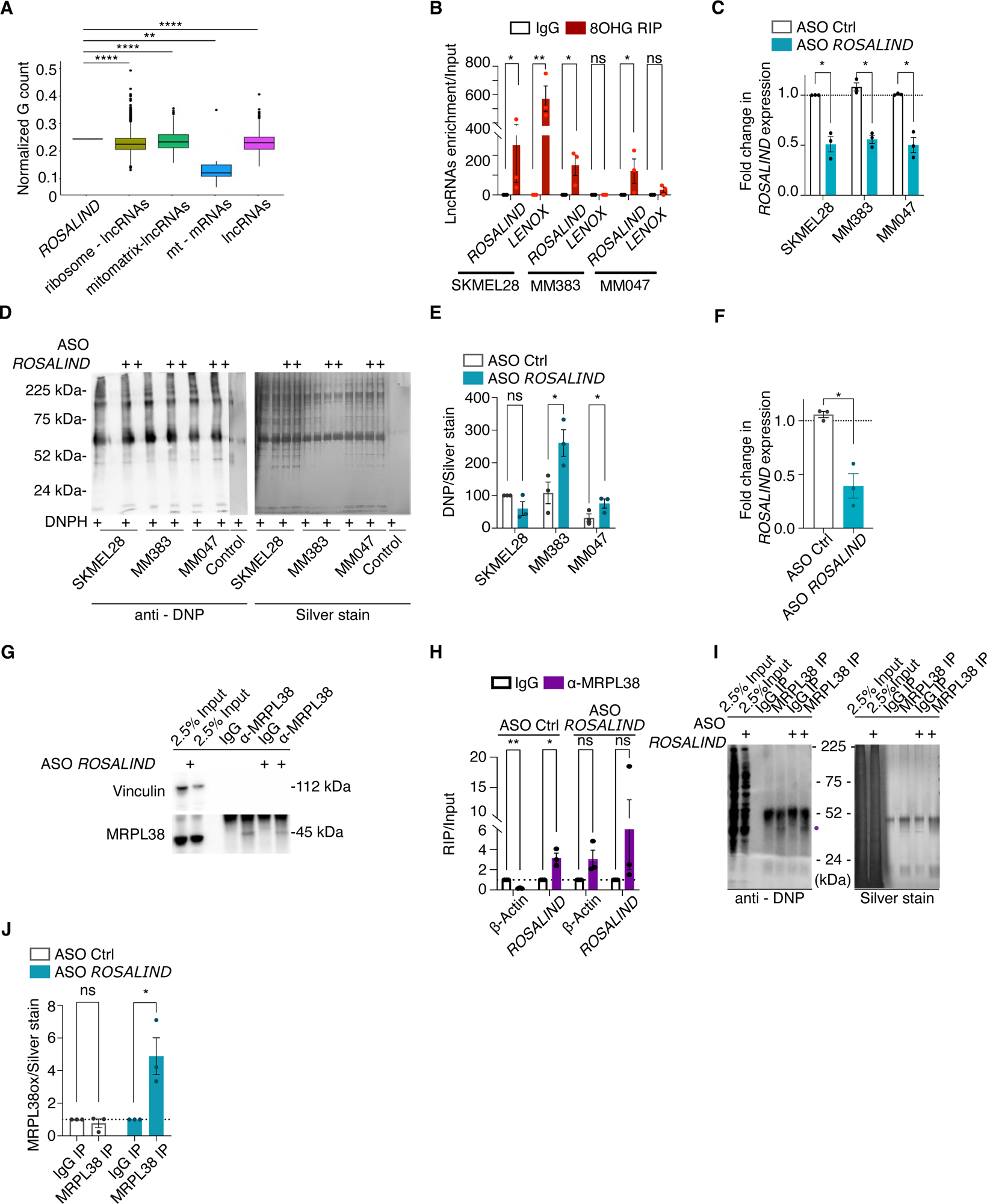
*ROSALIND* is a novel ROS buffering system. **A.** Percentage of G nucleotides normalised on transcript length. Significance was calculated by one-sample Wilkoxon Signed Ranked Test. 0**B.** *ROSALIND* enrichment over input in 8OHG RIP samples from SKMEL28, MM383 and MM047. Normal IgG were used as a control. Significance was calculated by multiple pared t-test. **C.** Fold changes in *ROSALIND* expression upon its KD in different melanoma lines. Significance was calculated by paired t-test. **D.** Western blot (left) and silver staining (right) of MM383, MM047 and MM099 mitoplast extracts upon inhibition of *ROSALIND*. Silver staining was used as a loading control. **E.** Densitometric analysis of DNP signal from the experiment in D. normalised on the silver staining. Paired t-test was used to calculate significance. **F.** Fold changes in *ROSALIND* expression upon its KD in PDX-derived MEL015 line. Significance was calculated by paired t-test. **G.** Efficiency of MRPL38 in the samples in F. as assessed by WB. Vinculin was used as a loading control. **H.** *ROSALIND* enrichment over input in MRPL38 RIP samples from PDX-derived MEL015 line. Normal IgG were used as a control. Significance was calculated by multiple pared t-test. **I.** Immunoprecipitation of MRPL38 upon inhibition of *ROSALIND* in MEL015 cells and detection of its oxidation status using anti-DNP antibody. Silver staining (right) was used as a loading control. **J.** Densitometric analysis of the western blot in I. Paired t-test was used to calculate statistical significance.

### *ROSALIND* inhibition makes melanoma cells more susceptible to immune cell killing

Defects in the ETC can lead to enhanced immune responses^23^, furthermore ROS production is associated with neoantigen generation and increased inflammation^24^. Having established that depletion of *ROSALIND* has cytotoxic effects due to oxidative damage of mitochondrial ribosomes and knowing that mitochondrial translation fidelity affects the performance of the ETC, we wondered whether *ROSALIND* KD increases the recognition of melanoma cells by the immune system. For this reason, performed *ROSALIND* KD in the PDX-derived melanoma cells Mel-015, derived from a model resistant to ICB^7^ and cocultured them with PBMCs activated with a T-cell stimulating cocktail (Figure 6A). Our data indicated that *ROSALIND* depletion increases melanoma cell killing by the immune system (Figure 6B-D).

**Figure 6A-M:**
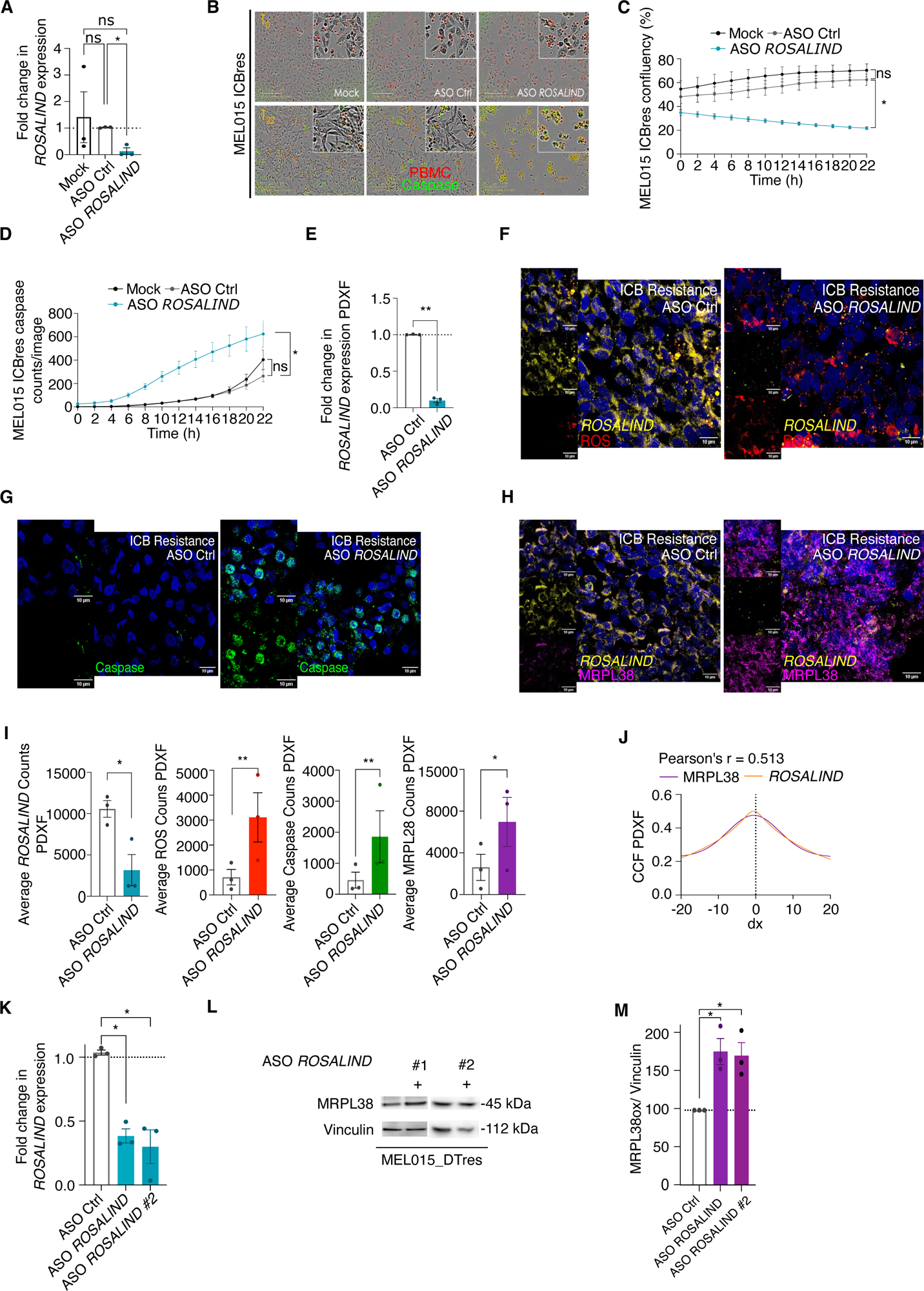
*ROSALIND* knock-down affects melanoma viability and immunogenicity. **A.** Fold changes in *ROSALIND* expression in MEL015_ICBres cells transfected with ASO *ROSALIND*, ASO control (ASO Ctrl) or not transfected (Mock). Significance was calculated by multiple paired t-test. **B.** Representative pictures of PBMCs (red) cocultured with MEL015_ICBres cells in A. Caspase-3+ cells are in green/yellow **C**. Analysis of MEL015_ICBres confluency (%) in A. Significance was calculated by two-way ANOVA mixed effect. **D**. Analysis of Caspase-3 counts in MEL015_ICBres transfected as indicated in A. Significance was calculated by two-way ANOVA mixed effect. **E.** Fold changes in *ROSALIND* expression in PDXFs derived from patient resistant to targeted or immune therapies, transfected with ASO *ROSALIND* or ASO control (ASO Ctrl). Significance was calculated by multiple paired t-test. **F.** Representative confocal images of the staining for ROS (red) and for *ROSALIND (*by FISH, yellow) of a PDXF model derived from humanised MEL015 resistant to immune therapy (Hu-Mel015), upon inhibition of *ROSALIND.* **G.** Representative confocal images of the PDXF described in G. stained by immunofluorescence for caspase (green). **H.** Representative confocal images of the PDXF described in G. stained by FISH for *ROSALIND* (yellow) and immunofluorescence for MRPL38 (purple). **I.** Quantification (average counts) of the stainings in Figure 6F-I. Significance was calculated by paired t-test. **J.** Colocalisation of *ROSALIND* with MRPL38 in the experiment shown in H., calculated using JaCoP Cross Correlation Function (CCF) with a pixel shift of δ = ±20. ρ indicates Pearson’s coefficient. **K.** Fold changes in *ROSALIND* expression upon its KD with two different ASO in the PDX-derived cell line MEL-015 DT resistant (MEL015_DTres). Significance was calculated by paired t-test. **L.** WB for MRPL38 in MEL015_DTres. Vinculin was used as a loading control. **M.** Densitometric quantification of the WB in L. Expression of MRPL28 was normalised on vinculin. Significance was calculated by paired t-test.

Lastly, to test the therapeutic potential of *ROSALIND* KD in 3D tumour models, we used ASO to inhibit it (Figure 6F) in PDX-derived tumour Fragments (PDXF) from melanoma models intrinsically resistant to immunotherapy (humanised with a human immune system Figure 6I-Q; Supplemental figure 6A-C) and to targeted therapy (Supplemental figure 6D-F). Silencing of *ROSALIND* (Figure 6F) and subsequent staining with MitoSOX Red detected elevated levels of ROS and caspase-3 *ex vivo* in preclinical patient-derived models (Figure 6F-G, I and Supplemental figure 6A-B and D-E). Furthermore, co-staining of MRPL38 and *ROSALIND* in these tumours revealed a strong colocalisation, validating the association of *ROSALIND* with the mitoribosome *ex vivo* (Figure 6H, J and Supplemental figure 6C and F). Surprisingly in these PDXFs, we observed by immunofluorescence a notable increase in MRPL38 levels in absence of localisation changes (Figure 6H; Supplemental figure 6C, F). This increase was validated with two different ASOs, by western blot on Mel-015 cells derived from PDX tumours that acquired resistance to targeted therapy (Figure 6K-M). It is therefore possible that upon inhibition of *ROSALIND*, MRPL38 is more oxidatively damaged and is thus more translated by the tumours to compensate for the subsequent mitochondrial translation inhibition. Taken together, our data suggest that targeting *ROSALIND* overcomes resistance to targeted and immune therapies.

## Discussion

The redox balance and the role of antioxidants in cancer is complex as their activity can tilt the equilibrium between cancer promotion and aging^25^. Collectively several lines of evidence support the idea that increased oxidative stress supports tumour initiation an metastatisation ^26^ however this effect is largely dependent on the genetic and metabolic context^10,26^. Furthermore, high ROS levels contribute to the efficacy of many anticancer drugs^12^ and increase cancer cell immunogenicity^22,27^. It is clear therefore that fine tuning of ROS levels can tilt the cell balance between life and death. This would explain while large randomized controlled clinical trials with antioxidants have failed to demonstrate benefit and sometimes even showed an increase in all-cause mortality^28^. In line with the finding that OXPHOS use is increased during tumour progression^4,8,9,17,29^ and therapy resistance, here we provided further evidence that cancer cells need a robust ROS scavenging system to persist and recur. Specifically, we described a previously unrecognized ROS buffering system based on the G-rich junk RNAs. The lncRNA *ROSALIND* is expressed in tissues with high oxidative metabolism such as the brain and the skin. Oxidative damage is also implicated in aging ^30^ and actually the antioxidant properties of RNAs are not new to the field of cosmetics. As a matter of fact, a company called Paula’s choice (https://www.paulaschoice.com/ingredient-dictionary/ingredient-sodium-rna.html) offers yeast RNA as an anti-aging skin solution. Here we unveil the molecular basis behind the RNA anti-aging effect and pave the way to the design of synthetic RNA molecules with antioxidant properties inspired by nature.

In cancer, *ROSALIND* gets anchored to the mitochondrial ribosomal proteins at the central protuberance where it protects the ribosomes and possibly the formation of the peptide bond from oxidative damage. The mitochondrial ribosomes diverge substantially from the bacterial ones as their RNA/protein ratio is inverted^31^. The newly acquired ribosomal proteins play a role in the synthesis of the highly hydrophobic proteins involved in the ETC that need to be inserted in the membrane co-translationally^31^. It follows that their preservation is vital for the correct assembly and functionality of the oxidative phosphorylation chain.

Compared to other antioxidants in the cell, *ROSALIND* has the advantage of not being incorporated in the housekeeping metabolic pathways of the cell and thus will escape most tumour suppressive feedback mechanisms in place. As many anti-cancer therapeutic modalities lead to ROS production ^12^, the reduction of cellular tolerance towards ROS would be an obvious adjuvant strategy. As a matter of fact, we have shown that inhibition of *ROSALIND* can overcome therapy resistance *in vitro* and *ex vivo* in preclinical models. Since redox balance is of outmost importance for healthy tissues, targeted inhibition in the tumour environment would be desirable. Considering its selective high expression in recurrent tumours, the inhibition of *ROSALIND* with ASOs would offer such an opportunity in multiple cancers. Lastly, our discovery suggests possibilities for the development of synthetic RNAs with antioxidant properties that could be used in industry for cosmetics or to prevent food deterioration.

## Material and methods

### Cell lines and transfection

SKMEL28 (from ATCC) were cultured in RPMI 1640 (Gibco BRL Invitrogen), supplemented with 10% FBS (Gibco BRL Invitrogen). The patient-derived low-passage MM cell lines (a gift from G.-E. Ghanem) were cultured in Ham’s F-10 (Gibco BRL Invitrogen), supplemented with 10% FBS (Gibco BRL Invitrogen). The PDX-derived cell lines MEL015_DTres and MEL015_ICBres were cultured in RPMI 1640 (Gibco BRL Invitrogen), supplemented with 10% FBS (Gibco BRL Invitrogen). To assess cell growth and viability, cells were stained with Trypan Blue (Invitrogen). All cell lines used are of human origin and were confirmed negative for mycoplasma before experimental use by using the MycoAlert Mycoplasma Detection Kit (Lonza) according to the the manufacturer’s instructions. For *ROSALIND* knock-down, 50 nM of a scramble ASO control or ASO against *ROSALIND* (for sequences refer to Table 1) was transfected using Lipofectamine LTX (Thermo Fisher Scientific) according to the manufacturer’s instructions. Cells were collected for RNA and/or protein extraction 48 h post transfection. For SOX10 knock-down, SKMEL28 cells were plated in 6 - well plates and transfected with Lipofectamine 2000 (Thermo Fisher Scientific) according to the manufacturer’s instructions with 50 nM non-targeting siRNA control (Horizon discovery) or with siSOX10 (ON-TARGET plus SMART-pool, Horizon discovery) 24 h after seeding. Cells were collected for RNA extraction 48 h post transfection. For *ROSALIND* overexpression, 150 - 300 ng of pcDNA3.1-LINC01918_201 or pcDNA3.1-empty vector were transfected using Lipofectamine LTX (Thermo Fisher Scientific) according to the manufacturer’s instructions. Cells were collected for RNA and/or protein extraction 48 h post transfection.

### ASO synthesis

Antisense oligonucleotides synthesis was performed on an Expedite 8909 DNA synthesiser (Applied Biosystems) by applying the phosphoramidite method. The ASOs, which are in the form of phosphorothioated LNA GapmeRs, were deprotected and cleaved from the solid support by treatment with AMA solution (1:1 mixture of ammonia 33% and methylamine 40%) for 2 h at 40°C. After gel filtration using an illustra NAP-25 column (Sephadex G-25 DNA Grade; GE Healthcare) and water as an eluent, the crude mixture was purified using a MonoQ HR 10/100 GL anion exchange column (GE Healthcare) with the following gradient system: 10 mM NaClO4 and 20 mM Tris-HCl in 15% CH_3_CN, pH 7.4 (A); 600 mM NaClO_4_ and 20 mM Tris-HCl in 15% CH3CN, pH 7.4 (B), 0-80% buffer B in 40 min, 2 ml/min. The low-pressure liquid chromatography system consisted of a Hitachi Primaide PM1110 HPLC pump, Mono-Q HR 10/100 GL column, and Hitachi Primaide PM1410 HPLC UV detector. The product-containing fractions were desalted on a NAP-25 column and lyophilized, and ASO sequences were analysed by mass spectrometry.

### Plasmid cloning

To clone *ROSALIND*, its cDNA fragment was amplified by PCR using as template an oligo-dT synthetized cDNA from a whole cell lysate of SKMEL28 cells. For this, *LINC01918_201* was amplified in two parts of approximately 400 nucleotides each: the first part was obtained using as forward primer GGCGGCGCCGCGGCCG, binding the beginning of the transcript and an intermediate reverse primer TCAGTGTGATTTAGAGGAATGAAATGCCAT. The second part was amplified using an intermediate forward primer GGGGCACAATGGCATTTCATTCCTCTAAAT, overlapping the intermediate reverse primer mentioned before and a primer binding at the end of the transcript AATTAAAGCATTCTTTAGGTAGGGTGAAAAC. The full-length transcript was then obtained using the forward and reverse primers mentioned above, binding the beginning and the end of the *LINC01918_201* sequence and using as template the two intermediate products. Subsequently, recognition sites for the restriction enzymes BamHI-HF (NEB) and EcoRI-HF (NEB) were added to the full-length transcript at the 5’ end and 3’ end respectively, using the primers CGC<u>GGATCC</u>GGCGGCGCCGCGGCCG as forward and CCG<u>GAATTC</u>AATTAAAGCATTCTTT as reverse. Both the full length *LINC01918_201* and the pcDNA3.1 empty vector (a kind gift from Prof. Chris Marine) were digested with BamHI-HF (NEB) and EcoRI-HF (NEB) restriction enzymes according to the manufacturer’s instructions. The linearized vector was then ligated to the digested PCR product using a quick ligase (NEB) according to the manufacturer’s instructions. The resulting vector was then used to transform DH5α competent cells (Thermo Fisher Scientific) and single clones were sent for Sanger DNA sequencing to verify the insertion and fidelity of *LINC01918_201* sequence in the pcDNA3.1 vector. *ROSALIND*sequence for overexpression (5’ to 3’) is: GGCGGCGCCGCGGCCGCGCAGCCTTTGTCTCGGGCCGCCGGGCGCGCGGGGCCCAGCGCAGGCAGGCCACGCGGGGACGGAGGAGGCAGAGAGGACGGGCTGTGATGGAGAAACGGGAGAGGAGCCGGCCAGGCGCCACCGTCACCACCATCACCAACCACCACCACCATCACCATCACTGACTGTCGCCCATCCCAGCTCCCCTGCCTCACCTCCCATCTCATATCCCTGTGCAGTGCCAACTTTATCAGATCATCTCTTTTTCACTGGGTACTACTTTGGAAAAGCGACAGAGTGCGCCAAGTCAGGAATAAGTACATCCGCTGTGGGACATTTATCTTCCCCTTTCTATTTAGCTACCTCAATTTTCTCATCAGGAAAGGGGGCACAATGGCATTTCATTCCTCTAAATCACACTGAGATGGCTGGGTTGCTTCCAGTTTTTCCAGTCCTGGCACTGGACGGTTATACATGCTTGGTGATGATGATGATAAAGTAATGTCTTCATTAAATTGGACAGTTGGGAGTTTTCAGACTTACTCTTCTTTGACTTCTGAGTGTCAGTATACCAGTAGCAAATAGAAACATAAGGAAAGTGAAACTTTGGCTAACATAATACCAAGAATAGATGCAAAACAGGCACTTTTGTAGGGACATCACCCAAGAGACCTGCAGGCAATGGTATTCATCTGCAGTGTAGAAAGCACCCCATCAGTGCCAAGTGGCTGTGGGAGCTCCTCAATCTGGATGCACAGTTTCCCGTGGGGTTTTCACCCTACCTAAAGAATGCTTTAATT

### Ascorbate Peroxidase (APEX)-proximity ligation assay and immunoaffinity purification

SKMEL28 cells (plated in 15 cm dishes, 8/condition) were transfected using lipofectamine 2000 (Thermo Fisher Scientific), according to the manufacturer’s instructions, with 10 μg of mito-V5-APEX (Addgene plasmid #72480) plasmid each. 24 h post-transfection the cells were treated with 30 μM Tigecycline (Selleck Chemicals), or PBS 1X (Sigma Aldrich) as a control, and were left for additional 24 h. Before harvest, all cells were treated with 500 μM biotin-phenol (Iris Biotech) and incubated for 30 min, 37°C, 5% CO_2_. After incubation with the APEX substrate, the cells were treated with 1 mM H_2_O_2_ (Sigma Aldrich) for 1 min, at Room Temperature (RT), agitating in the dark, to catalyse the biotinylation of all proteins residing in the mitochondrial matrix and in close proximity to the APEX. The cells were afterwards washed several times in quencher solution [10 mM sodium ascorbate (VWR), 10 mM sodium azide (VWR), 5 mM Trolox (Sigma Aldrich)] to neutralise the remaining free radicals, followed by washes in PBS 1X (Sigma Aldrich). Subsequently the cells were UV-crosslinked using 0.4 J/cm^2^ (UVP Crosslinker Analytik Jena), collected in PBS 1X (Sigma Aldrich) and lysed for 20 min on ice in 1 ml lysis buffer [Tris-HCl pH 8.0 20 mM, NaCl 200 mM, MgCl_2_ 2.5 mM, Triton-X 1% (Sigma Aldrich), DTT 1 mM (Sigma Aldrich), Halt Protease and Phosphatase Inhibitor Cocktail 1X (Fisher Scientific), SUPERase·In 60 U/ml (Fisher Scientific) in DEPC-H_2_O (VWR)] supplemented with quencher solution. Protein concentration was measured using the QubiT Protein Assay (Thermo Fisher Scientific) according to the manufacturer’s instructions. Before proceeding with the pull-down, 60 μg of each sample were kept as protein and RNA input. The rest of the sample was divided into equal amounts of protein (1.5 mg of protein/pull-down) and incubated with 80 μl of pierce streptavidin magnetic beads (Thermo Fisher Scientific), O/N, 4°C, rotating. The following day, the bead-biotinylated protein complexes were washed twice in lysis buffer, thrice in DEPC-treated H_2_O (VWR) and twice more in lysis buffer. Each sample was then divided as follows: 1/3 of the sample was used for protein elution in order to validate by western blot analysis the efficiency and accuracy of the IP and 2/3 of the sample were used for RNA extraction and subsequent library preparation. Protein elution was performed by adding 30 μl 6X Laemmli on dry beads followed by boiling the sample at 95°C, 15 min, in constant shaking. For RNA extraction, 500 μl of TRIzol (Thermo Fisher Scientific) were added on dry beads and RNA was extracted following the manufacturer’s instructions. RNA samples were prepared for sequencing using the TruSeq Stranded Total RNA kit (Illumina) according to the manufacturer’s instructions. Libraries were sequenced using the HiSeq4000. These data have been deposited in GEO database with accession number GSE249588.

### Mitoplast isolation

Mitochondria were isolated using the Mitochondria Isolation Kit for Cultured Cells (Thermo Fisher Scientific) according to the manufacturer’s instructions. Briefly, cells grown in 15 cm plates were scraped and collected in falcons using PBS 1X (Sigma Aldrich). The cells were subsequently lysed using the reagents supplied by the kit, supplemented with SUPERase·In 60 U/ml (Fisher Scientific) and Halt Protease and Phosphatase Inhibitor Cocktail 1X (Fisher Scientific). Isolated mitochondria were further processed with RNase A containing buffer [10 μg/ml RNase A (Sigma Aldrich), 10 mM HEPES pH 7.2, Halt Protease and Phosphatase Inhibitor Cocktail 1X (Fisher Scientific)] to eliminate contaminating cytosolic RNA for 20 min on ice followed by 10 min RT incubation, washed twice in mitoplast isolation buffer [250 nM Mannitol (Sigma Aldrich), 5 mM HEPES pH 7.2 (Sigma Aldrich), 0.5 mM EGTA (Sigma Aldrich), 1 mg/ml bovine serum albumin (BSA) (Sigma Aldrich) supplemented with SUPERase·In 60 U/ml (Fisher Scientific) and Halt Protease and Phosphatase Inhibitor Cocktail 1X (Fisher Scientific) in DEPC-H_2_O (VWR)] and were ultimately lysed to collect RNA and/or protein for downstream applications. For proteinase K experiments, purified mitochondria were resuspended in 1X sucrose buffer [HEPES 10 mM (Sigma Aldrich), sucrose 280 mM (Sigma Aldrich), EGTA 1 mM pH 7.4 (Sigma Aldrich)] supplemented with 100 µg/ml proteinase K (Sigma Aldrich) and SUPERase·In 60 U/ml (Fisher Scientific) for 30 min, 4°C, rotating. Mitoplasts were then pelleted at 9.000 rcf for 10 min and then resuspended in 1X sucrose buffer supplemented with 20 nM PMSF (Sigma-Aldrich) and SUPERase·In 60 U/ml (Fisher Scientific) and pelleted again at 9.000 rcf for 10 min.

### Mitoplast-ribosome profile

SKMEL28 cells were plated in 30 15 cm plates at 70% confluency on the day of the experiment. Before harvest, cells were treated with 100 μg/ml chloramphenicol (Sigma Aldrich) for 30 min, 37°C, 5% CO_2_ to immobilise mitochondrial ribosomes. The cells were subsequently fractionated as described before, and mitoplasts were isolated with proteinase K treatment (Sigma Aldrich). Isolated mitoplasts were lysed for 20 min on ice in 300 μl lysis buffer [20 mM Tris-HCl pH 7.4, 10 mM NaCl, 3 mM MgCl_2_, supplemented with 1.2% Triton-X (Sigma Aldrich), 0.2 M Sucrose (Sigma Aldrich), 100 μg/ml chloramphenicol (Sigma Aldrich), Halt Protease and Phosphatase Inhibitor Cocktail 1X (Fisher Scientific), SUPERase·In 60 U/ml (Fisher Scientific)]. After removal of the mitoplast membranes by centrifuging at 17.000 rcf for 15 min at 4°C, mitoplast extracts were loaded on linear, 10% - 30% sucrose gradients prepared in buffer G (20 mM Tris-HCl, 100 mM KCl, 10 mM MgCl_2_, 1 mM DTT (Sigma Aldrich), 100 μg/ml chloramphenicol (Sigma Aldrich)] and ultracentrifuged using an SW41-Ti rotor (Beckman Coulter) at 37.000 rpm, 4°C for 170 min. The fractions were obtained with a Biological LP System (Bio-Rad). A total of 18 (500 μl) fractions were collected from each sample and were subsequently pooled together accordingly, to obtain the mitochondrial subunits 28S and 39S and 55S monosome.

### RAP-MS

To identify the protein targets of *ROSALIND,* 50 μl of protein A Dynabeads (Invitrogen) were coupled to 500 pmol of pooled biotinylated probes (IDT) against *ROSALIND* and incubated O/N at 4°C, rotating. As a negative control we used pooled biotinylated probes against *PCA3* (IDT), a lncRNA not expressed in melanoma samples (for sequences refer to Table 1). Tumour samples that acquired resistance to targeted therapy (Dabrafenib-Trametinib [DT]), or tumour samples that had not received DT, were rinsed in PBS 1X (Sigma Aldrich) and chopped using sterile scalpels into small pieces that could fit through a 1000 μl tip. The tumour pieces were then placed in a 50 ml falcon and incubated for 45 min at 37°C in RPMI 1640 (Gibco BRL Invitrogen) supplemented with 3% FBS (Gibco BRL Invitrogen), 0.3 µg/ml liberase (Sigma Aldrich) and 1X DNase I (Sigma Aldrich). Dissociated cells were spined down at 300 rcf for 5 min, washed twice with PBS 1X (Sigma Aldrich) and finally were passed through a cell strainer. Before proceeding to downstream applications, red blood cells had to be removed from the tumour-dissociated cell population. For this purpose, cells were incubated and inverted for 10 min at RT with ammonium chloride lyse [NH_4_Cl 155 mM (Sigma Aldrich), NaHCO_3_ 10 mM (Sigma Aldrich), EDTA 0.1 mM in ddH_2_0 (Fisher Scientific)] and were then spun down for 10 min at 250 rcf at 4°C. The supernatant was discarded, since it contained lysed red blood cells, and the remainder cells were washed twice in PBS 1X (Sigma Aldrich). These cells were subsequently lysed in 1 ml of “pull-out buffer” [20 mM Tris-HCl pH 7.5, 200 mM NaCl, 2.5 mM MgCl_2_, 0.05% NP-40 (Thermo Fisher Scientific), 1 mM DTT (Sigma Aldrich), SUPERase·In 60 U/ml (Fisher Scientific) and Halt Protease and Phosphatase Inhibitor Cocktail 1X (Fisher Scientific)] and incubated with the probes-beads mixture for 3 h, at 4°C, rotating. After this incubation, the beads were washed six times in lysis buffer and were then divided to obtain RNA and protein extracts, using the elution methods explained before. For mass spectrometry, the samples were run shortly on SDS-PAGE gels (Thermo Fisher Scientific) and stained with Coomassie brilliant blue (Biorad) to help in their excision. The excised pieces were washed thrice in 500 μl ddH_2_O (Fisher Scientific) and were then dried in a speed-vac. 50-150 μl of 10 mM DTT (Sigma Aldrich) diluted in 50 mM NH_4_HCO_3_ (ammonium bicarbonate; Sigma Aldrich) was added to the dried bands and incubated at 56°C for 45 min. After the incubation period the supernatant was removed and the samples were further incubated with 150 μl of 51.5 mM IAA (Merck Millipore) diluted in 50 mM NH_4_HCO_3_ (Sigma Aldrich) at RT for 30 min, in the dark. The supernatant was removed and excised pieces were washed thrice in 500 μl of 50 mM NH_4_HCO_3_ (Sigma Aldrich). The pieces were subsequently dried again using a speed-vac. For in-gel digestion of proteins, 50 mM NH_4_HCO_3_ (Sigma Aldrich), 0.01 % ProteaseMAX (Promega), 10% acetonitrile (Sigma Aldrich) and 3 ng/μl trypsin (Promega) were added to the dried pieces and incubated on ice for at least 40 min. Subsequently, the samples were overlayed with equal volumes of 50 mM NH_4_HCO_3_ (Sigma Aldrich), 0.01 % ProteaseMAX (Promega) and 10% acetonitrile (Sigma Aldrich) and incubated O/N at 37°C. The following day, the supernatant was collected in a new 1,5 ml tubes and the remaining peptides were extracted from the gel pieces with a series of incubations. First, 150 μl of ddH_2_O (Fisher Scientific) were added and incubated for 30 min at 37°C. Second, 150 μl of 60% acetonitrile (Sigma Aldrich) and 0.1% TFA (Sigma Aldrich) were incubated for another 30 min at 37°C. After each of these incubations, the supernatant was retrieved and pooled with the initial extract. The final extract was then adjusted to 1% TFA (Sigma Aldrich) and the samples were centrifuged for 10 min at 15.000 rcf. The supernatant was collected and dried in a speed-vac for 7 h. The obtained dried peptides were then processed using Pierce C18 columns (Thermo Fisher Scientific) according to the manufacturer’s instructions, to clean them from contaminants that could interfere with spectra quality. For UPLC separation, an Ultimate 3000 UPLC system (Thermo Fisher Scientific) equipped with an Acclaim PepMap100 pre-column (C18, particle size 3 μm, pore size 100 Å, diameter 75 μm, length 20 mm, Thermo Fisher Scientific) and a C18 PepMap analytical column (particle size 2 μm, pore size 100 Å, diameter 50 μm, length 150 mm, Thermo Fisher Scientific) using a 40 min linear gradient (300 nL/min) of 0-4% buffer B (80% ACN, 0.08% FA) for 3 min, 4-10% B for 12 min, 10-35% for 20 min, 35-65% for 5 min, 65-95% for 1 min, 95% for 10 min, 95-5% for 1 min, and 5% for 10 min, was used. The Orbitrap Elite Hybrid Ion Trap-Orbitrap mass spectrometer (Thermo Fisher Scientific) was operated in positive ion mode (nanospray voltage 1.8 kV, source temperature 275°C) in data-dependent acquisition mode with a survey MS scan at a resolution of 60.000 (FWHM at *m*/*z* 200) for a mass range of *m*/*z* 375-1.500 for precursor ions, followed by MS/MS scans of the top 20 most intense peaks above a threshold count of 500 with charge +2 or higher. Low resolution CID MS/MS spectra were acquired in rapid CID scan mode using a normalised collision energy of 35 eV and an isolation window of *m*/*z* 2.0. Dynamic exclusion was set to 10 s. Raw MS data were converted into .mgf files and subjected to database searching using Proteome Discoverer (version 1.4) software (Thermo Fisher Scientific). Peptides were identified by MASCOT (version 2.2.06, Matrix Science) using UniProt UniProt Homo sapiens (204906 sequences, 09/06/2022) as database. The following MASCOT search parameters were used: Trypsin/P, carbamidomethyl (C) as fixed modification, oxidation (M) as variable modification, two missed cleavages, peptide tolerance 10 ppm for MS and 0.5 Da for MS/MS. Progenesis software (version 4.1.4832.42146, Nonlinear Dynamics) was used for relative quantification of peptides. RNA interactomics and input data were analysed separately. Proteome Discoverer software was used for peptide annotation and peptide validation, the latter using the Percolator node (max delta Cn = 0; max rank = 1). Only peptides with a q value < 0.01 in any of the conditions were taken into account. RNA interactomics and input data were analysed separately. Scaffold software was used to validate MS/MS based peptide and protein identifications. Peptide and protein identifications were accepted to achieve both an FDR less than 1,0% and proteins contained at least 1 identified peptide. The protein candidate that was pursued for this work was identified with at least two different peptides in each pull-down sample and zero peptides in the negative control samples.

### RIP

For RIP, 1-2 mg of protein or 1-2 μg of RNA was used. All proteins or nucleotide modifications were immunoprecipitated using 5 μg of antibody, and same amount of an IgG (Merck Millipore) serving as a negative control, which was incubated with the lysate O/N at 4°C, rotating. The following day, each sample was incubated with 60 μl of protein A Dynabeads (Invitrogen) for 3 h at 4°C, rotating.

### RIP/RAP efficiency

Relative expression of the genes of interest from the RIP and RAP experiments was calculated applying the ΔΔCt method. Briefly, the Ct value (i.e. the number of cycles where the sample’s reaction curve was detected to intersect with the threshold line during the qPCR reaction) of each gene of interest was subtracted from the Ct value of the input, thus obtaining a ΔCt. Subsequently, a ΔCt was also calculated for each gene for the IgG or control probe sample, using the same principle. Ultimately, the ΔCt of the RIP/RAP was subtracted from the ΔCt of the control thus obtaining a ΔΔCt value. For the fold change calculation, the following formula was applied: fold change = 2^(-ΔΔCt)^. For the 8OHG RIP, the overall efficiency of RNA retrieval was also calculated. Total RNA counts from each 8OHG RIP were normalised on Anti-Rabbit Alexa-488 antibody that served as a baseline control. The same was applied to each IgG RIP. Finally, each normalised 8OHG RIP was normalised on its IgG control RIP. The obtained values depict the amount of total RNA that was pulled down upon 8OHG RIP. Generated graphs were made through GraphPad Prism v9.5. Paired t-test was used to calculate statistical significance.

### Incucyte-coculture experiments

For Incucyte experiments (cell viability assay), cell lines were transfected using Lipofectamine LTX (Thermo Fisher Scientific) with an ASO against *ROSALIND* or a scramble ASO control or were given only lipofectamine (Mock), according to the manufacturer’s instructions. The cells were then monitored by the IncuCyte ZOOM system (Essen BioScience) and four images per well were taken every 2 h. Cell death was assessed by adding the IncuCyte Caspase 3/7 Green Apoptosis Assay Reagent (1:5000 dilution) in the cell media (Essen BioScience). Cell confluency and caspase positivity were measured and analysed using the IncuCyte ZOOM software. The same technology was used for the coculture experiments using PBMCs (Precision for medicine) at 1:5 ratio. Knock-down of *ROSALIND* was performed as previously described in immunotherapy resistant (intrinsic) MEL015 cells. 48 h post transfection, pre-activated PBMCs were added to the cells and were followed for up to 22 h. For T cell activation, freshly thawed PBMCs were cultured in RPMI 1640 (Gibco BRL Invitrogen), supplemented with 10% FBS (Gibco BRL Invitrogen), 1% penicillin/streptomycin (Sigma-Aldrich), 3 μg/ml of human ⍺-CD3 (Thermo Fisher Scientific), 5 μg/ml of human ⍺-CD28 (Thermo Fisher Scientific) and 200 ng of IL-2 (Immunotools) for 48 h before being introduced to the melanoma cells. PBMCs were labelled with 1:1000 Incucyte® Nuclight Rapid Red Dye for Live-Cell Nuclear Labeling Incucyte (Sartorius) while cell death was measured by the IncuCyte Caspase 3/7 Green Apoptosis Assay Reagent as explained before. Generated graphs for % confluency and caspase counts/image were made through GraphPad Prism v9.5. Two-way ANOVA mixed effects was used to calculate statistical significance.

### MitoSOX staining *ex vivo* in Patient-derived Tumour Fragments (PDTF)

The cutaneous melanoma PDX models are part of the Trace collection (https://gbiomed.kuleuven.be/english/research/50488876/54502087/Trace). huMEL015-αPD1, MEL082, and MEL058 mouse models were used for the PDTFs (derived from male-drug naïve BRAFmut, female NRASmut and male-drug naïve BRAFmut patients respectively), the two first being intrinsically resistant to immunotherapy and the last intrinsically resistant to targeted therapy. Each tumour was cut using a sterile scalpel to a size small enough to fit in a 96-well plate. The tumours were plated on a coated 96-well plate containing tumour media [DMEM (Gibco BRL Invitrogen), 1 mM C_3_H_3_NaO_3_, 1X MEM (Gibco BRL Invitrogen), 2 mM L-glutamine (Thermo Fisher Scientific), 10% FBS (Gibco BRL Invitrogen), 1% penicillin/streptomycin (Sigma-Aldrich)] supplemented with 1 mg/ml collagen I (Corning) and 1.1% sodium bicarbonate (Sigma Aldrich). 24 h after plating, the tumours were transfected using Lipofectamine 2000 (Thermo Fisher Scientific) with an ASO against *ROSALIND* or a scramble ASO control, according to the manufacturer’s instructions. 48 h post transfection the tumours were incubated with 100 μM MitoSOX Red (Thermo Fisher Scientific) for 1 h, at 37°C, 5% CO_2_. Before downstream applications, the tumours were cut in two pieces: one piece would serve for immunofluorescence (IF) and the other piece would be lysed in Trizol (Thermo Fisher Scientific) to obtain RNA and validate the knock-down efficiency. The tumour pieces used for IF were fixed in 4% formol (VWR) for 45 min at RT and were then placed in cryovials at -25°C for 45 min to freeze. Multiple 10 μm slices were cut from each tumour piece using a cryostat that was set at -25°C. The tumours were then permeabilised in 70% EtOH for 2 h at RT and then blocked with blocking buffer [1% BSA (Sigma Aldrich), 0.2% Triton-X (Sigma Aldrich), 10% goat serum (Abcam), PBS 1X (Sigma Aldrich)] for 30 min at RT. They were then incubated with FISH probes against *ROSALIND* (9 pooled probes tagged with a fluorophore excited at 488 nm, IDT) and/or antibodies against MRPL38 (1:500) (PA5-118074, Thermo Fisher Scientific) and Caspase 3/7 (1:300) [Cleaved Caspase-3 (Asp175) (5A1E) (Cell Signaling)], O/N at 37°C in a humidified dark chamber. The following day the tumours were incubated with appropriate secondary antibodies for 45 min at RT and were then imaged using the Nikon C2 confocal microscope. For image acquisition, the same confocal settings were applied to all images stemming from all mouse models.

### Carbonyl assay

SKMEL28, MM383 and MM047 cell lines were plated in 2 15 cm dishes and transfected using Lipofectamine LTX (Thermo Fisher Scientific) with an ASO against *ROSALIND* or a scramble ASO control according to the manufacturer’s instructions. 48 h post transfection the cells were collected, and the mitoplast was isolated as explained previously. The protein carbonyl assay kit (western blot) (Abcam ab178020) was used to assess the oxidation status of mitochondrial proteins according to the manufacturer’s instructions. Briefly, mitoplast extracts were lysed in 1X extraction buffer for 20 min on ice and membranes were discarded by centrifuging at 17.000 rcf, 15 min, at 4°C. Each supernatant was denatured with 6% SDS and was subsequently split into two equal parts: one part was derivatized by the addition of 2,4-Dinitrophenylhydrazine (DNPH) for 15 min at RT and the other part served as a negative control of the reaction. After the incubation period, both parts were neutralised and were run on SDS-PAGE gels (Thermo Fisher Scientific) without boiling. Two partner gels were loaded for this experiment: the first gel would serve for the detection of protein carbonyl groups that are created by the oxidation of proteins and the second gel would serve as a loading control, by performing a silver staining (Thermo Fisher Scientific) according to the manufacturer’s instructions. For western blotting, membranes were incubated with anti-DNP antibody (1:5000 dilution) O/N at 4°C and then with HRP-secondary antibody (1:5000 dilution) for 1 h at RT. Membranes were developed using ECL (Radiance or Radiance Plus, Sopachem) and images were acquired in an Azure 600 Imaging System (azure biosystems). Relative protein levels were quantified using ImageJ and statistical significance was calculated by performing paired t-test.

### RNA extraction

For RNA extraction of ribosomal fractions, each fraction was digested at 37°C for 90 min in a mix of 100 μg/ml proteinase K (Sigma-Aldrich) and 1% SDS (Sigma Aldrich). Phenol acid chloroform (5:1; Sigma-Aldrich) and 10 mM NaCl were subsequently added to the samples. The samples were then centrifuged at 16.000 rcf for 5 min at 4°C. The upper aqueous phase was transferred to a new tube, and 1 ml of isopropanol (Fisher Chemical) was added. Samples were placed at −80°C O/N to precipitate RNA. The following day, the samples were centrifuged at 16.000 rcf for 40 min at 4°C. The supernatant was discarded, and the RNA pellet was washed using 500 μl of 70% EtOH, then centrifuged again at 16.000 rcf for 5 min at 4°C. Pellets were air-dried for a maximum of 5 min and resuspended in DEPC-treated water (VWR). For all the other experiments, RNA was extracted using TRIzol (Thermo Fisher Scientific) and subjected to DNase treatment (Invitrogen) according to the manufacturer’s instructions.

### RT and qPCR

RNA was reverse transcribed using the High-Capacity complementary DNA Reverse Transcription Kit (Thermo Fisher Scientific) according to the manufacturer’s instructions on a Veriti 96-well thermal cycler (Thermo Fisher Scientific). Gene expression was measured by qPCR on a QuantStudio 5 system (Thermo Fisher Scientific) and normalised by applying the ΔΔCt method using GAPDH or 18S as reference genes (for mito-polysome experiments) or on the average of GAPDH, UBC, and ACTB for other experiments. Sequences of the primers are indicated in Table 1. For the evaluation of lncRNA polyadenylation, total RNA was reverse transcribed with the High-Capacity complementary DNA Reverse Transcription Kit (Thermo Fisher Scientific) using either random hexamers (Thermo Fisher Scientific) or an Oligo(dT)12-18 Primer (Thermo Fisher Scientific) according to the manufacturer’s instructions, on a Veriti 96-well thermal cycler (Thermo Fisher Scientific). Polyadenylation was assessed by qPCR on a QuantStudio 5 system (Thermo Fisher Scientific) and normalised by applying the ΔΔCt method using transcript-specific primers. GAPDH, UBC, and ACTB were used again as reference genes. Generated graphs were made through GraphPad Prism v9.5. (Un)paired t-test was used to calculate statistical significance.

### RNA sequencing and data analysis

Samples were prepared for sequencing using TruSeq Stranded Total RNA kit (Illumina) according to the the manufacturer’s instructions. Libraries were sequenced on an Illumina NextSeq and HiSeq4000 according to the the manufacturer’s instructions, generating PE high output configuration cycles: (R1: 100) − (I1: 6) − (I2: 0) − (R2: 100) and (R1: 50) − (I1: 6) − (I2: 0) − (R2: 0). Differential gene expression analysis was performed as previously described^4^.

### Protein extraction and western blot

Proteins were extracted by resuspending the cellular pellet in RIPA buffer [150 mM NaCl, 50 mM Tris-HCl pH 8, 1% Nonidet P40 (Thermo Fisher Scientific), 0.5% Sodium Deoxycholate (Sigma Aldrich), 1 mM EDTA (Sigma Aldrich)] supplemented with Halt Protease and Phosphatase Inhibitor Cocktail 1X (Fisher Scientific)]. Western blotting experiments were performed using the following primary antibodies at the indicated dilution: vinculin (V9131, clone VIN-1; Sigma-Aldrich, 1:5.000), XRN2 (A301-103A; Bethyl, 1:2.000), ATP5A (7H10BD4F9; Abcam, 1:1.000), NDUFS3 (17D95; Abcam, 1:1.000), TOM20 (F-10; Santa Cruz, 1:500), MRPL38 (PA5-118074; Thermo Fisher Scientific, 1:1.000), Histone H3 (D1H2; Cell signaling, 1:1.000), Catalase (ab217793; Abcam, 1:1.000), Puromycin (12D10; Santa Cruz, 1:10.000), Beta-actin (13E5; Cell signaling, 1:5.000). The following HRP-linked secondary antibodies were used: HRP-conjugated Streptavidin (N100; Thermo Fisher Scientific, 1:1.000), ⍺-mouse IgG (NA931-1ML; Sigma-Aldrich, 1:5.000) and ⍺-rabbit IgG (NA934-1ML; Sigma-Aldrich, 1:5.000). Relative protein levels were quantified using the Image Studio Lite 5.2 software and plotted using GraphPad Prism v9.5. Paired t-tests was used to calculate statistical significance.

### Puromycin labelling of nascent peptides *in vitro* (SUNsET)

To monitor changes in cytosolic and mitochondrial translation cells were labelled with puromycin (Invivogen) and SUNsET was performed as previously described^32^. In brief, cells were plated at a 40% - 50% confluency and the next day either knock-down or overexpression of *ROSALIND* was performed as described earlier. 48 h post-transfection the cells were washed twice in PBS 1X (Sigma Aldrich) and were pulsed with 10 μg/ml of puromycin (Invivogen) diluted in media, for 15 min, 37°C, 5% CO_2_. The media was refreshed, and the cells were placed again at 37°C, 5% CO_2_ for additional 30 min (“chase”), to ensure that puromycin would also penetrate the mitochondrial membranes. Since puromycin is a tRNA analogue, it gets incorporated into nascent peptides but prevents the ribosome from peptide elongation, causing their release from the ribosome. Puromycin can be fatal *in vitro* if used at very high concentrations and for a prolonged period, but if used appropriately it can be an indicator of mRNA translational rates both in the cytosol and in mitochondria. To monitor the two translational machinery rates separately, mitoplasts were fractionated as described before and puromycin incorporation was assessed by western blot using an anti-puromycin antibody, as previously described.

### Immunofluorescence

For immunofluorescence, cells were typically grown in 6-well plates after placing round coverslips in the bottom of the wells (VWR), or in PhenoPlates 96-well (Perkin Elmer). Cells were fixed in 3.7% formaldehyde (Sigma Aldrich) for 20 min at RT and permeabilised in permeabilization buffer [1% BSA (Sigma Aldrich) and 0.2% Triton X-100 (Sigma Aldrich) in PBS 1X (Sigma Aldrich)] for 10 min on ice. Blocking was performed in 10% goat serum (Abcam) diluted in permeabilization buffer, for 30 min at RT. Primary antibody incubation was performed at RT for 1 h, in blocking buffer. For the detection of MRPL38, 1:500 dilution of the antibody was used (PA5-118074, Thermo Fisher Scientific), for P32 1:1.000 (A302-863A, Bethyl laboratories), for TOM20 1:500 (clone F-10, sc-17764, Santa Cruz) and for Caspase 3/7 1:300 [Cleaved Caspase-3 (Asp175) (5A1E) (Cell Signaling)]. Secondary antibody incubation was performed for 45 min at RT in the dark, in blocking buffer. As secondary antibodies, ⍺-rabbit or ⍺-mouse AlexaFluor-488 and AlexaFluor-647 (Life Technologies, 1:1.000) were used. Nuclei were stained with 1:1.000 DAPI (Sigma Aldrich) diluted in PBS 1X (Sigma Aldrich) for 15 min at RT before mounting the slides using the ProLong™ Glass Antifade mounting medium (Thermo Fisher Scientific). Slides were then imaged on a Nikon C2 confocal microscope while the 96 wells were imaged by the high content screening microscope Operetta CLS+ Twister (Perkin Elmer). Images were analysed using Image J and Harmony softwares (Perkin Elmer) respectively. Average counts were measured from at least three focal fields from each of three biological replicates. Statistical significance was calculated by unpaired t test. Colocalization analysis was performed using the Fiji Image J plug-in JACoP; Van Steensel on z-stacks of confocal images. Evaluation of colocalization levels was measured with cross correlation function (CCF) with a pixel shift of δ = ±20.

### Fluorescent *in situ* hybridization

For detection of *ROSALIND*, we used a pool of 9 FISH probes (IDT) (Table 1) baring FAM, which were designed using the Stellaris probe designer software (Biosearch Technologies). Cells were grown in 6-well plates after placing round coverslips in the bottom of the wells (VWR), or in PhenoPlates 96-well (Perkin Elmer), fixed in 3.7% formaldehyde (Sigma Aldrich) for 10 min at RT and permeabilised in 70% EtOH for at least 1 h at RT. Cells were then washed twice in wash buffer [2X sodium lauryl sulfate (Sigma Aldrich), 10% formamide (Sigma Aldrich)] and hybridized with 250 nM of pooled probes diluted in hybridization buffer [2X sodium lauryl sulfate (Sigma Aldrich), 10% formamide (Sigma Aldrich) and 10% dextran (Sigma Aldrich)], overnight at 37°C inside a humidified chamber. Cells were washed twice in wash buffer for 30 minutes at 37°C inside a humidified chamber and thrice with PBS 1X (Sigma Aldrich). Nuclei were stained with 1:1.000 DAPI (Sigma Aldrich) diluted in PBS 1X (Sigma Aldrich) for 15 minutes at RT before mounting the slides using the ProLong™ Glass Antifade mounting medium (Thermo Fisher Scientific). Slides were then imaged on a Nikon C2 confocal microscope while the 96 wells were imaged by the high content screening microscope Operetta CLS+ Twister (Perkin Elmer). Images were analysed using Image J and Harmony softwares (Perkin Elmer) respectively. Average counts were measured from at least three focal fields from each of three biological replicates. Statistical significance was calculated by unpaired t test. Colocalisation analysis was performed using the Fiji Image J plug-in JACoP; Van Steensel on z-stacks of confocal images. Evaluation of colocalisation levels was measured with cross correlation function (CCF) with a pixel shift of δ = ±20.

### Colony formation assay

MM383, MM047, MM099, SKMEL28 and MEL015_DTres cells were plated in 6-well or 12-well plates at 30% - 50% confluency and cultured for approximately 1 week. On the desired day, the cells were washed twice in PBS 1X (Sigma Aldrich) and subsequently fixed and stained for 20 min using a solution containing 1% crystal violet (Sigma Aldrich) and 35% methanol (Thermo Fisher Scientific) diluted in dH_2_O. Knock-down or overexpression of *ROSALIND* was performed as described previously. For N-Acetyl-L-Cysteine (NAC) (Sigma Aldrich) experiments, NAC was used at a concentration of 2.5 mM diluted in media, 24 h post-transfection. Surface occupancy was assessed using ImageJ. Paired t-tests were used to calculate statistical significance.

### MitoTracker staining and mito-shape properties evaluation

To assess the properties of mitochondrial shape, MEL015_DTres cells were cultured in 6-well plates containing round coverslips in the bottom (VWR) and were transfected with an ASO targeting *ROSALIND* or a scramble ASO control as previously described. 48 h post-transfection the cells were stained with 200 nM MitoTracker CMXRos Red (Thermo Fisher Scientific) for 30 min, at 37°C, 5% CO_2_. They were then fixed using 3.7% formaldehyde (Sigma Aldrich) for 20 min at RT and the slides were mounted using the ProLong™ Glass Antifade mounting medium (Thermo Fisher Scientific). Slides were subsequently imaged on a Nikon C2 confocal microscope. Image J was used to quantify the properties of mitochondrial shape and network upon knock-down of *ROSALIND* as has been demonstrated before^8,33,34^. Briefly, 12 z-stack sections/condition of a biological triplicate containing multiple cells each, were each merged into a single image by generating a maximum intensity composite which was pre-processed using the “subtract background” (radium 1um), “sigma filter plus” (radius 0.1 μm, 2.0 sigma), “enhance local contrast/CLAHE” (block size 64, slope 2.0), “gamma correction” (0.8), and tubeness (sigma 0.361) commands on Image J. The “adaptive threshold” plugin was used to identify mitochondria in the images, which were subsequently processed using the command “despeckle”. The resulting binary images were used as the input for the “analyse particles” command, measuring for area, perimeter, and shape descriptors. Form factor (FF) was derived as the inverse of the circularity^35^. For network connectivity analysis, the “skeletonize 2D/3D” command was applied to produce a skeleton map of mitochondria inside each cell and the “analyse skeleton” command was used to calculate the number of networks. The results were visualised using GraphPad Prism. In the generated graphs, each dot represents a processed z-stack (as explained above) containing multiple cells, that have been averaged for each of their mitochondrial properties. Unpaired t-test was used to calculate statistical significance.

### Measurement of ROS production by flow cytometry

For assessment of ROS levels, cells were grown in 6-well plates at 30%-50% confluency and the following day *ROSALIND* knock-down was performed as previously described. 48 h post-transfection the cells were washed twice in PBS 1X (Sigma Aldrich) and incubated with 250 nM MitoSOX Red (Thermo Fisher Scientific) diluted in media without addition of FBS for 30 min, in the dark, at 37°C, 5% CO_2_. The cells were subsequently washed thrice in PBS 1X (Sigma Aldrich) in the dark, resuspended in 300 μl F10 media (Gibco BRL Invitrogen) containing 5% FBS (Gibco BRL Invitrogen) and 2% EDTA and passed through a sterile filter-top FACS tube (Fisher Scientific). The samples were analysed on a BD FACSymphony flow cytometer. For ROS rescue experiments, 24 h post-transfection the cells were treated with 2.5 mM N-Acetyl-L-Cysteine (Sigma Aldrich) and 48 h post-treatment were stained with MitoSOX Red as described above.

### Analysis of oxygen consumption rate in living cells (Seahorse)

Oxygen consumption rates (OCR) were measured using a Seahorse XFp Analyzer (Agilent). 2.000 cells were plated per well of a XFp miniplate at a final volume of 80 μl. The following day, the cells were transfected with an ASO targeting *ROSALIND* or a scramble ASO control as previously described. 48 h post-transfection the media was replaced with RPMI 1640 pH 7.4 (Gibco BRL Invitrogen) supplemented with 10 mM glucose (Agilent), 1 mM sodium pyruvate (Agilent) and 2 mM L-glutamine (Agilent) and the cells were placed in a non-CO_2_ incubator set at 37°C for approximately 1 h before starting the assay. After the incubation period, the cells were sequentially exposed to 2 μM oligomycin (Sigma Aldrich), 20 μM FCCP (Cayman Chemical-Sanbio) and 0.5 μM antimycin A (Sigma Aldrich) and were followed for 73 min. The obtained OCR (pmoles/min) values were plotted using GraphPad Prism v9.5 and two-way Anova mixed effect was used to calculate statistical significance. To obtain separate graphs for basal respiration, ATP production, maximal respiration and spare capacity, the average of the three measurements before the addition of a new compound was calculated. Paired t-test was used to calculate statistical significance.

### Single-cell RNA sequencing analysis (scRNAseq)

Single-cell sequencing data were generated from human melanoma patients after a single dose of ICB treatment^20^. The data were then processed using the “Seurat” package in R. Subsequently, the R packages “ggpubr” and “ggplot” were used to create graphs for gene expression, using the aforementioned dataset^36^.

### GSEA from RNA seq data

Raw counts were downloaded from the GEO database project “Single-cell analysis of gene expression variation and phenotype switching in melanoma” (data accessible at NCBI GEO database, accession GSE134432). Count-based differential expression analysis was done with the R-based (R Foundation for Statistical Computing) Bioconductor package “DESeq”^37^. The resulting list of differentially expressed genes was used for GSEA.

From the scRNA seq data mentioned above, “FindMakers” was used to determine the differentially expressed genes between cells that express *ROSALIND* as compared to cells that do not and these were subsequently used to perform GSEA.

Geneset enrichment analysis^38^ was performed using GSEA 4.2.2. on the fold change of significantly differentially expressed genes (*P* value < 0.05). Plots visualising GSEA data were made using GraphPad Prism v9.5 and only genesets with nominal *P* value < 0.05 and FDR q value < 0.1 are shown.

### CPC2

The coding potential of the *LINC01918_201* was predicted by the CPC2 algorithm^41^. Graphpad prism was used to visualize the results.

### Statistical analysis

All statistical analyses performed are indicated on each experimental section. *P* values are represented as follows: *P* > 0.05, not significant (ns); *P* < 0.05, *; *P* < 0.01, **; *P* < 0.001, ***; *P* < 0.0001, ****. All statistical analyses were performed on GraphPad Prism v9.5 or in R.

## Author’s contributions

VK performed most of the wet lab experiments and assembled figures. YV performed all the bioinformatic analysis. ELJ performed Incucyte analysis in Figure 4 and supplementary figure 3. ED prepared PDTF and PDX-derived cell lines. EG provided ASO. AM and RD designed and interpreted mass spectrometry experiments. VK and EL designed the study. EL interpreted the data and wrote the manuscript. All authors read and edited the manuscript.

## Acknowledgements

We would like to thank the Light Microscopy and Imaging Network - LiMoNe (VIB-KULeuven) and Guy Schepers and Frédérick Coosemans (Rega Institute) for the excellent technical support with microscopy and with ASO synthesis respectively. We would like to thank Dr. G. Gambi and Dr. R. Vendramin for the useful discussion and assistance with the mitochondrial measurements. The present study was funded by a KU Leuven C1 grant, a Belgian federation for Cancer grant (FAF-F/2018/1184). E. Leucci was funded by the Melanoma Research Alliance Amanda and Jonathan Eilian young investigator award. V. Katopodi is a recipient of a FWO – Flanders research organization PhD fellowship (1S47519N). Y. Verheyden is FWO – Flanders research organization PhD fellowship (1SC5122N). S. Cinque is a recipient of FWO – Flanders research organization PhD fellowship (1SD1620N).

Sequencing data have been deposited to GEO database with accession number GSE249588. Proteomics data have been deposited to ProteomeXchange consortium with the following accession number PXD047963.

---

**Figure S1A-L:**
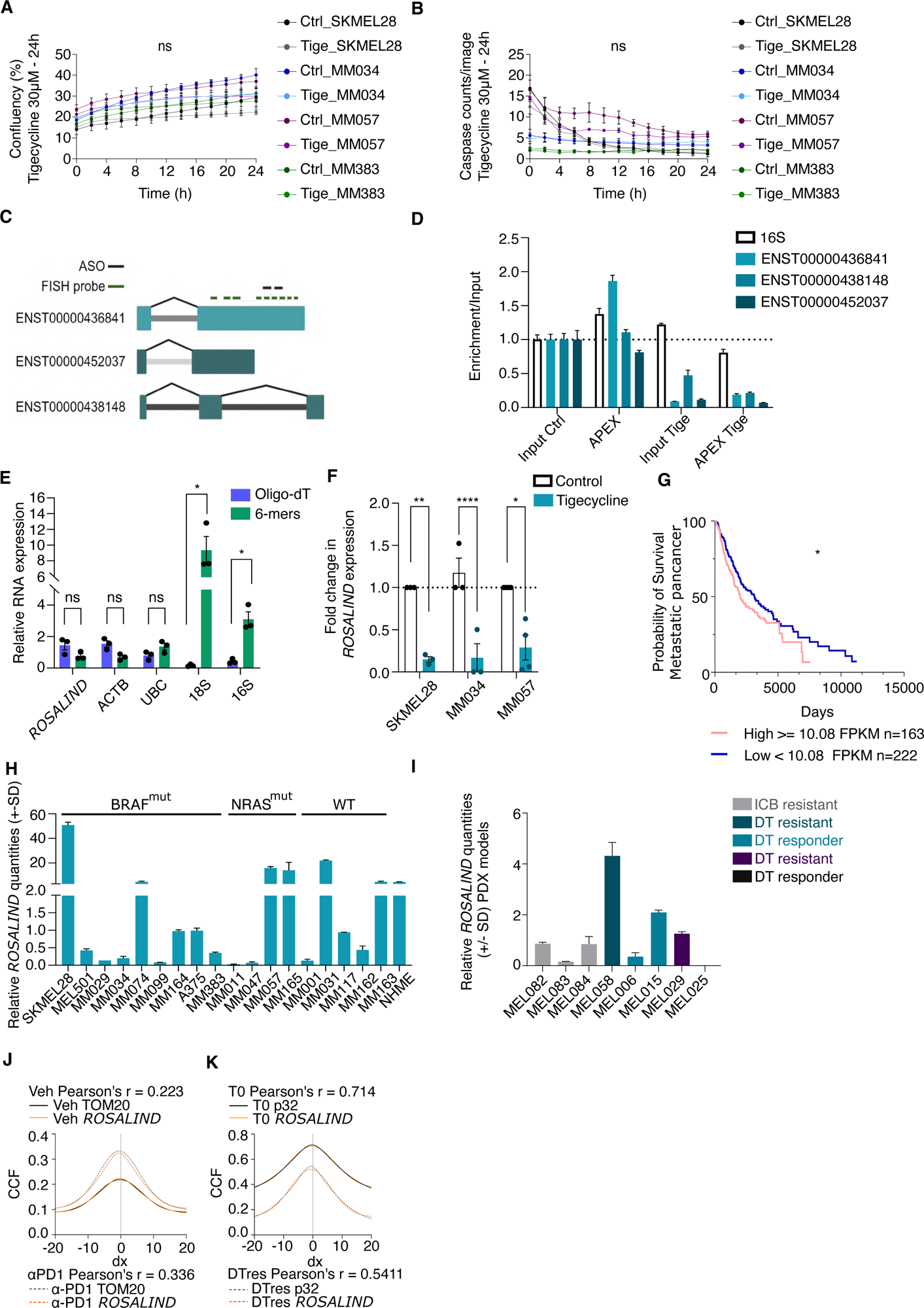
*ROSALIND* expression is sensitive to mitochondrial translation inhibition and correlates with its binding partner MRPL38. Analysis of SKMEL28, MM034, MM057, MM0383 confluency (%), as measured by Incucyte, upon addition of 30μM Tigecycline. Two-way ANOVA mixed effect was used to calculate statistics **B.** Analysis of SKMEL28, MM034, MM057, MM0383 Caspase-3 counts upon addition of 30μM Tigecycline. Two-way ANOVA mixed effect was used to calculate statistics **C.** Graphic representation of *ROSALIND* isoforms and binding of FISH probes and ASO. **D.** Enrichment over input of different *ROSALIND* isoforms in the APEX-seq samples as assessed by qPCR. **E.** Relative expression of polyadenylated and non-polyadenylated *ROSALIND* isoforms as assessed by RT-qPCR with random hexamers or oligo-dT. *16S* and *18S* were used as a control of non-polyadenylated transcripts and *actin B* as a control of polyadenylation. Significance was calculated by multiple paired t-test. **F.** Fold changes in *ROSALIND expression* in SKMEL28, MM034 and MM057 melanoma cells upon treatment with 30μM tigecycline for 24h. Significance was calculated by paired t-test. **G.** Correlation between *ROSALIND* expression and patients’ survival by Kaplan-Meier plots in the metastatic patients of the PanCancer. Significance was calculated by log-rank (Mantel-Cox) test. **H.** Relative *ROSALIND* expression in a panel of melanoma cell lines, representative of different phenotypic states and carrying different driver mutations, and Normal Human Melanocytes. **I.** Relative *ROSALIND* expression in a panel of untreated melanoma PDX models derived from drug-naïve and drug-resistant patients. **J.** Colocalisation of *ROSALIND* and TOM20 in PDX-derived samples on treatment with α-PD1, calculated using JaCoP Cross Correlation Function (CCF) with a pixel shift of δ = ±20. ρ indicates Pearson’s coefficient. **K.** Colocalisation of *ROSALIND* and p32 in PDX-derived samples on treatment with targeted therapy, calculated using JaCoP Cross Correlation Function (CCF) with a pixel shift of δ = ±20. ρ indicates Pearson’s coefficient.

**Figure S2A-K:**
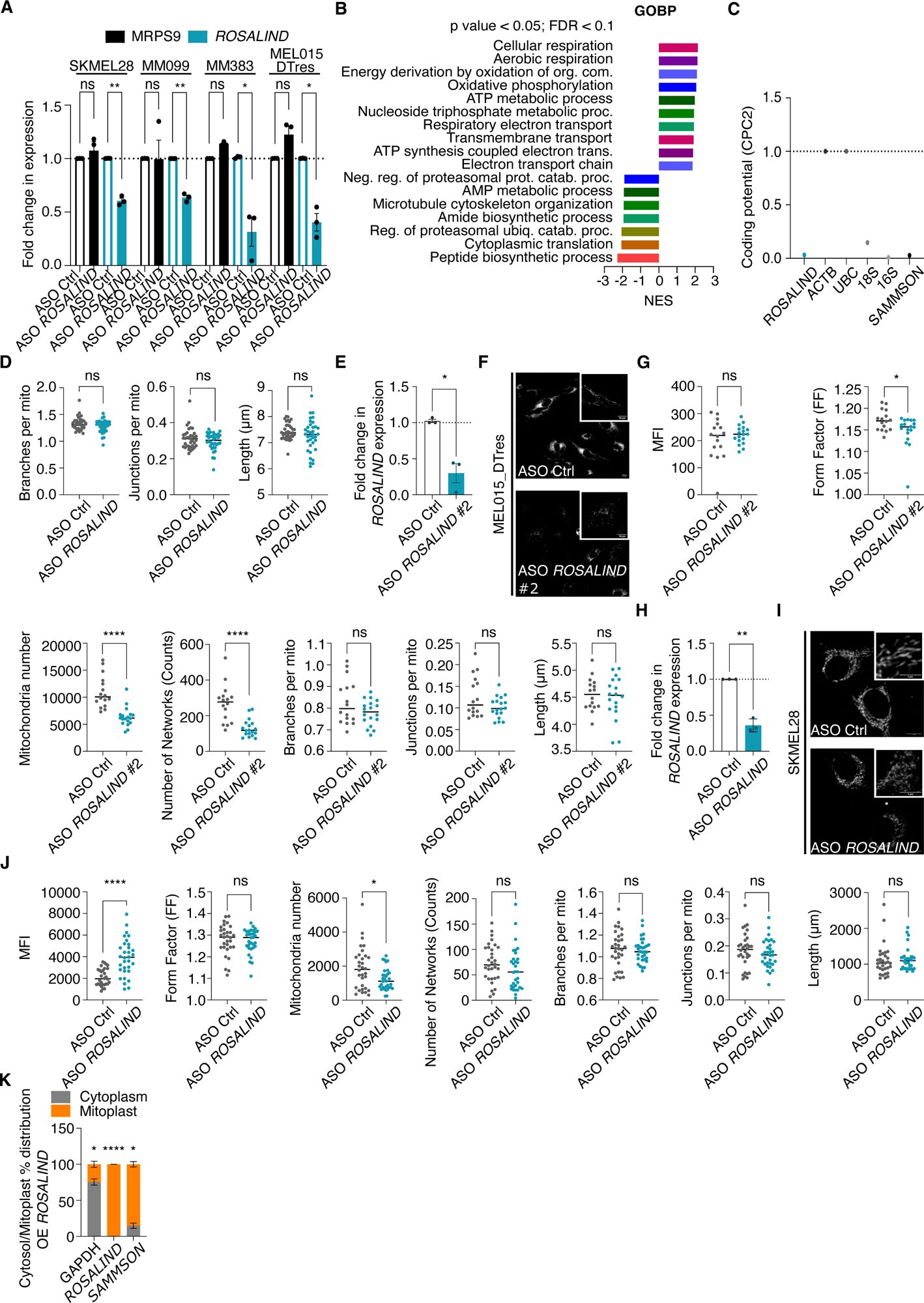
*ROSALIND* affects mitochondrial functions. **A.** Fold changes in *ROSALIND* and MRPS9 expression in SKMEL28, MM099, MM383 and MEL015_DTres cells transfected with ASO *ROSALIND* or ASO control (ASO Ctrl). Significance was calculated by multiple paired t-test. **B.** GSEA on the differentially expressed proteome of the PDX-derived tumours used for *ROSALIND* puIl-down. Threshold values FDR<0.1 p<0.05. **C.** In silico prediction of *ROSALIND* protein-coding potential. Actin B and UBC have been used as a positive control, *SAMMSON* and rRNAs as a negative control. **D.** Quantification of mitochondrial properties from the images shown in Figure 3O where *ROSALIND* was inhibited in MEL015_DTres cells with MiNa workflow on ImageJ. Each dot represents an image acquired from the confocal with multiple cells, that have been averaged for each of the mitochondrial properties. Images used for these graphs are derived from a biological triplicate. Significance was calculated by unpaired t-test. **E.** Fold changes in *ROSALIND* expression in MEL015_DTres cells transfected with ASO *ROSALIND #2* or ASO control (ASO Ctrl). Significance was calculated by paired t-test. **F.** Representative confocal images from Mitotracker Red stained MEL015_DTres cells upon inhibition of *ROSALIND* with ASO #2. **G.** Quantification of mitochondrial properties from the images shown in F. where *ROSALIND* was inhibited with ASO #2 in MEL015_DTres cells with MiNa workflow on ImageJ. Each dot represents an image acquired from the confocal with multiple cells, that have been averaged for each of the mitochondrial properties. Images used for these graphs are derived from a biological triplicate. Significance was calculated by unpaired t-test. **H.** Fold changes in *ROSALIND* expression in SKMEL28 cells transfected with ASO *ROSALIND* or ASO control (ASO Ctrl). Significance was calculated by paired t-test. **I.** Representative confocal images from Mitotracker Red stained SKMEL28 upon inhibition of *ROSALIND*. **J.** Quantification of mitochondrial properties from the images shown in I. where *ROSALIND* was inhibited with ASO #2 in MEL015_DTres cells with MiNa workflow on ImageJ. Each dot represents an image acquired from the confocal with multiple cells, that have been averaged for each of the mitochondrial properties. Images used for these graphs are derived from a biological triplicate. Significance was calculated by unpaired t-test. **K.** *ROSALIND* cytosolic/mitochondrial ratios (in percentage) as assessed by RT-qPCR on mitoplast extracts of SKMEL28 cells upon its overexpression. *SAMMSON* and *GAPDH* were used as a mitochondrial and cytosolic marker respectively. Significance was calculated by multiple paired t-test.

**Figure S3A-Q:**
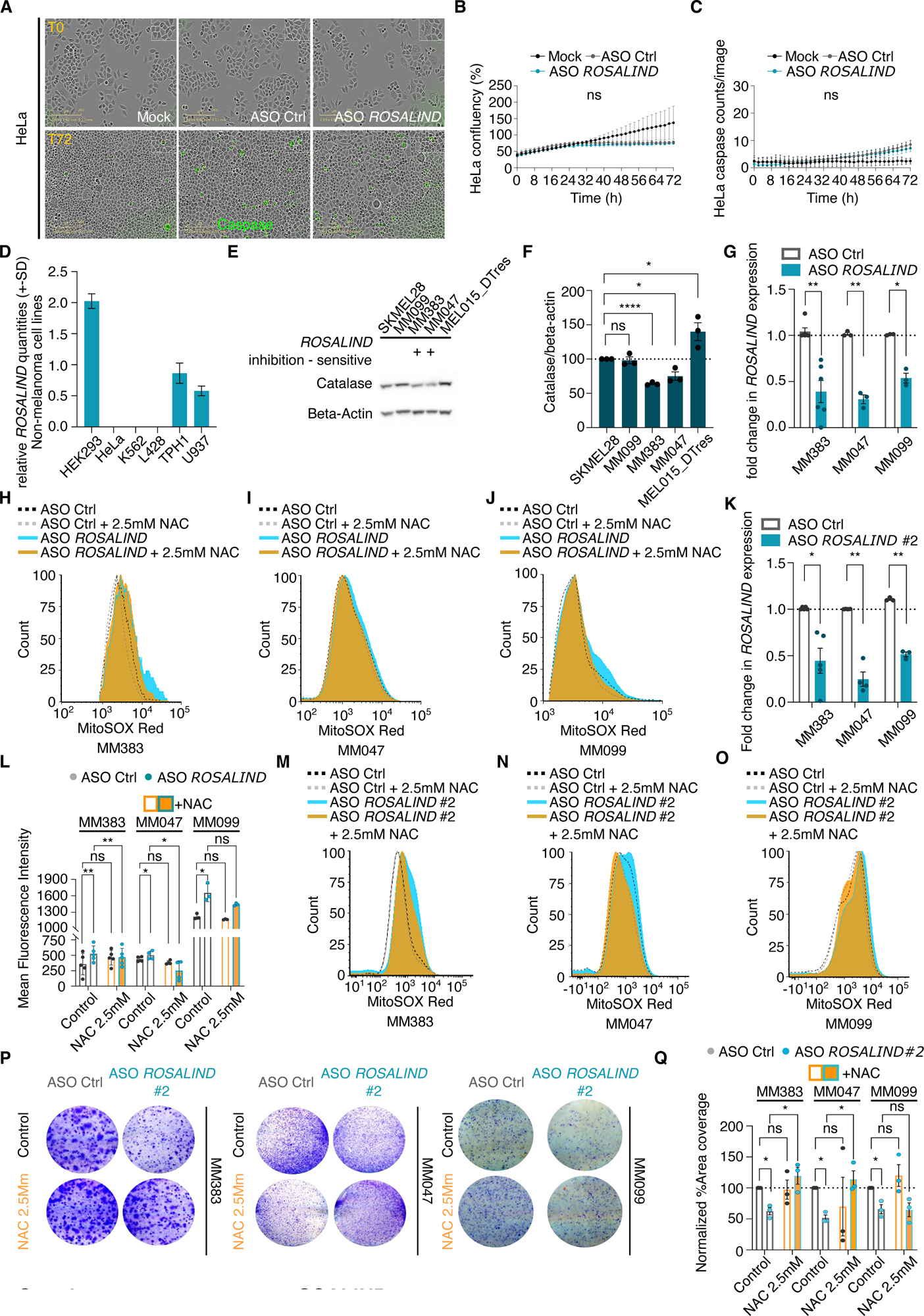
*ROSALIND* affects melanoma cell viability. **A.** Representative Incucyte pictures from HeLa cells at T0 and T72 h upon inhibition of *ROSALIND.* Caspase-3+ cells are in green. **B**. Analysis of HeLa confluency (%) transfected with ASO *ROSALIND*, ASO control (ASO Ctrl) or not transfected (Mock). Two-way ANOVA mixed effect was used to calculate statistics. **C.** Analysis of Caspase-3 counts in HeLa upon transfected as indicated in S3B. Two-way ANOVA mixed effect was used to calculate statistics. **D.** Relative *ROSALIND* expression in HeLa cells as compared to other cancer cell lines. **E.** Western blot for catalase in several melanoma cell lines sensitive and resistant to *ROSALIND* inhibition. Beta-actin was used as a loading control. **F.** Densitometric analysis of the western blot in E. Significance was calculated by unpaired t-test was used to calculate statistical. **G.** Fold change in *ROSALIND* expression upon its inhibition with ASO#2 in MM383, MM047 and MM099. Significance was calculated by paired t-test. **H.** Representative FACS histogram of MM383 cells stained with MitoSOX Red, corresponding to the experiment shown in Figure 4O **I.** Representative FACS histogram of MM047 cells stained with MitoSOX Red, corresponding to the experiment shown in Figure 4O. **J.** Representative FACS histogram of MM099 cells stained with MitoSOX Red, corresponding to the experiment shown in Figure 4O. **K.** Fold change in *ROSALIND* expression upon its inhibition with ASO#2 in MM383, MM047 and MM099. Significance was calculated by paired t-test. **L.** Evaluation of ROS production by MM383, MM047 and MM099 upon *ROSALIND* ASO #2 transfection with (orange bars) and without (grey bars) NAC buffering, as measured by MitoSOX staining (mean fluorescent intensity) and flow cytometry analysis. Significance was calculated by two-way ANOVA. **M.** Representative FACS histogram of MM383 cells stained with MitoSOX Red, corresponding to the experiment shown in L. **N.** Representative FACS histogram of MM047 cells stained with MitoSOX Red, corresponding to the experiment shown in L. **O.** Representative FACS histogram of MM099 cells stained with MitoSOX Red, corresponding to the experiment shown in L. **P.** Colony assays of MM383, MM047 and MM099 upon transfection of *ROSALIND* ASO#2 in presence or absence of NAC buffering. **Q.** Quantification of colony assays in P expressed as percentage of area covered normalised to the Control sample (treated only with ASO Ctrl, without NAC). Significance was calculated by two-way ANOVA.

**Figure S4A-D:**
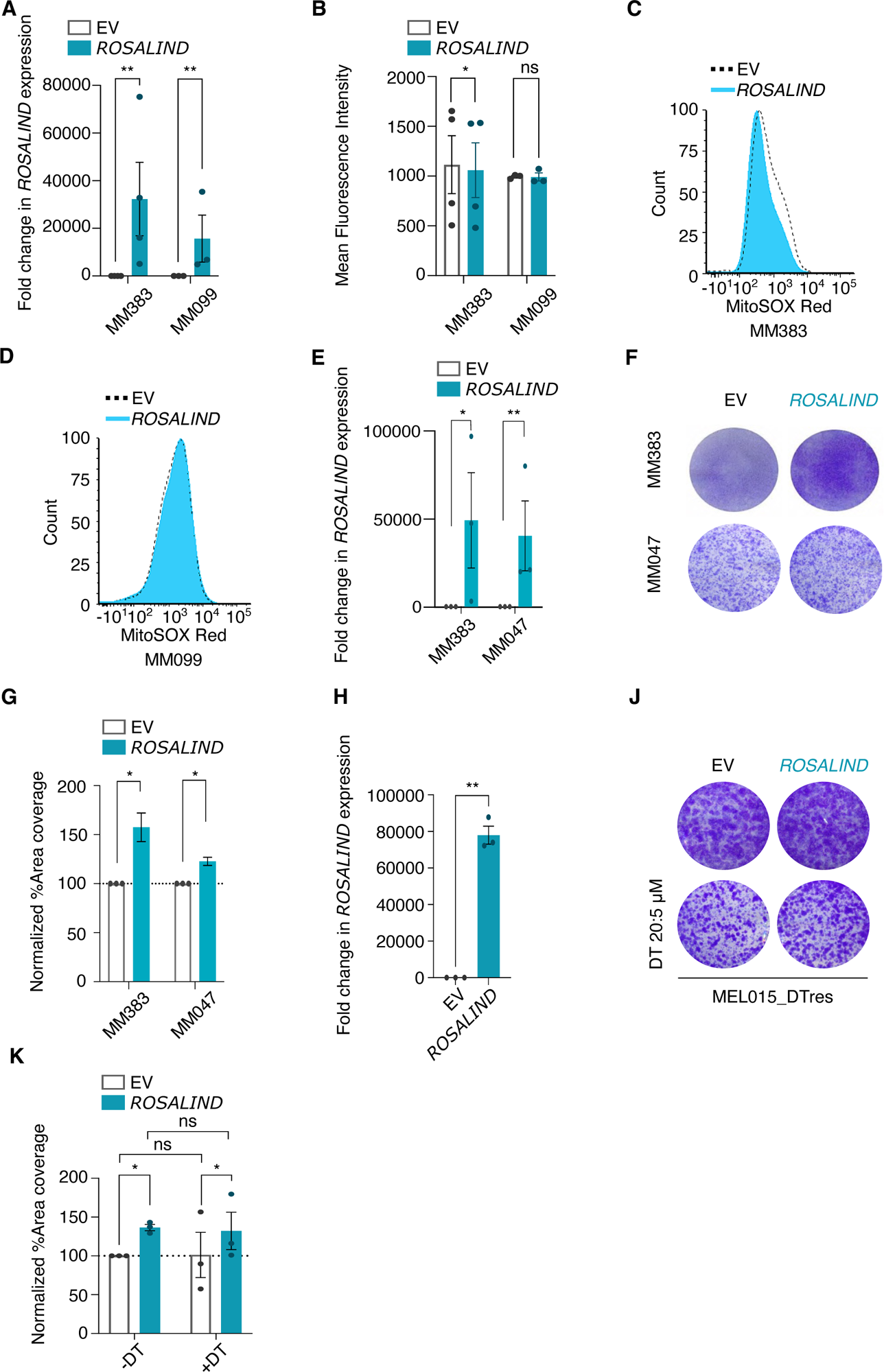
*ROSALIND* overexpression improves melanoma cell viability. **A.** Fold changes in *ROSALIND* expression upon its ectopic expression in MM383 and MM099 cells. Significance was calculated by multiple paired t-test. **B.** Evaluation of ROS production by MM383 and MM099 upon *ROSALIND* overexpression. Significance was calculated by two-way ANOVA. **C. and D.** Representative FACS histogram of MM383 (C.) and MM099 cells (D.) stained with MitoSOX Red, corresponding to the experiment shown in C. **E.** Fold changes in *ROSALIND* expression upon its ectopic expression in MM383, MM047 and MM099 cells. Significance was calculated by multiple paired t-test. **F.** Colony assays of MM383 and MM047 upon *ROSALIND* overexpression **G.** Quantification of colony assays in F. expressed as percentage of area covered normalised to the control sample transfected with an empty vector (EV). Significance was calculated by multiple paired t-test. **H.** Fold changes in *ROSALIND* expression upon its ectopic expression in MEL015_DTres. Significance was calculated by paired t-test. **I.** Colony assays of MEL015_DTres upon *ROSALIND* overexpression combined with targeted therapy treatment (DT). **J.** Quantification of colony assays in F. expressed as percentage of area covered normalised to the control sample transfected with an empty vector (EV). Significance was calculated by two-ways ANOVA.

**Figure S5A-D:**
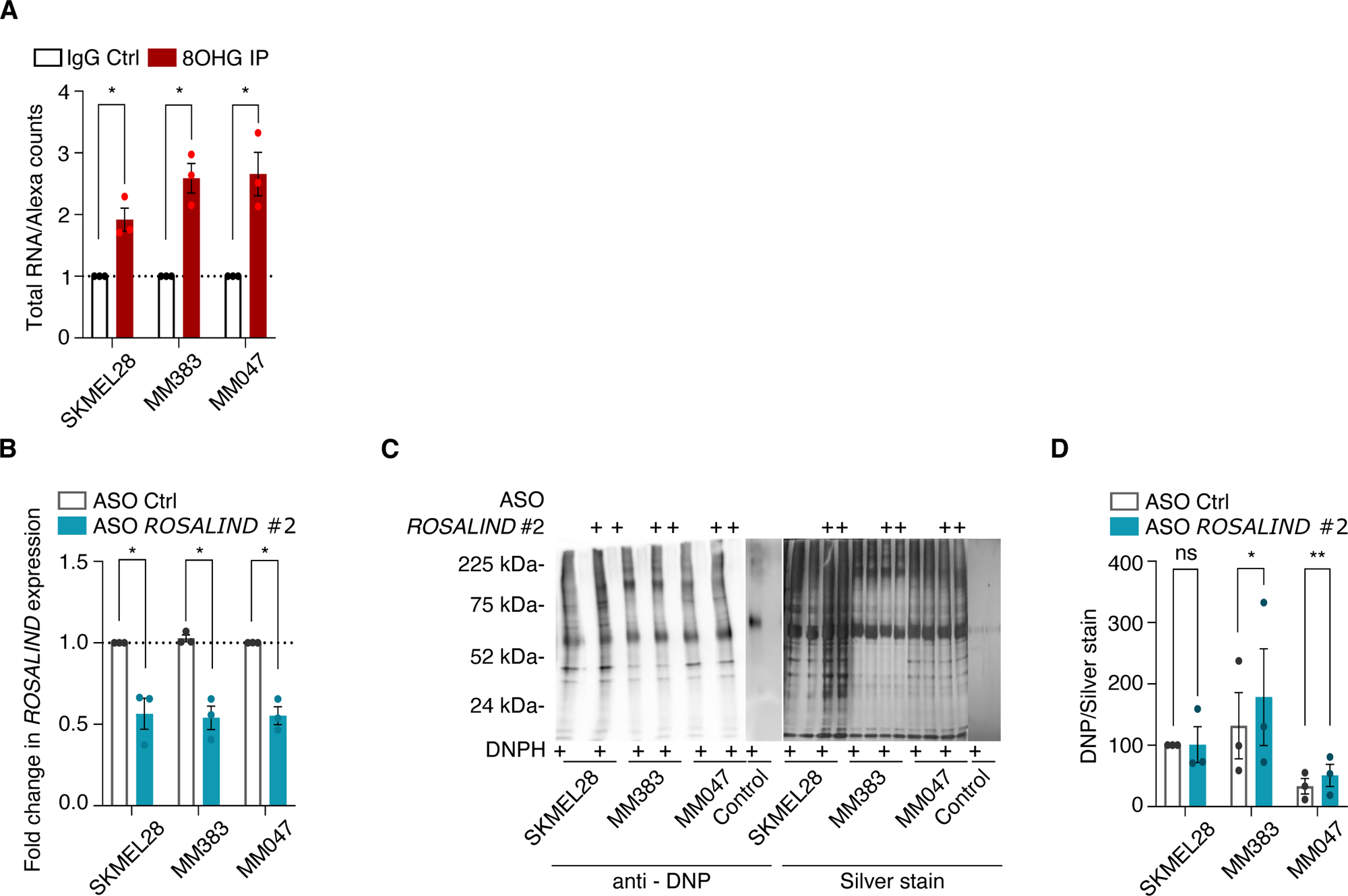
*ROSALIND* scavenges ROS. **A.** Efficiency of the 8OHG immunoprecipitation shown in Figure 5B. Total RNA counts from each 8OHG IP were normalised on Anti-Rabbit Alexa-488 antibody that served as control and subsequently normalised on IgG control IP. Paired t-test was used to calculate significance. **B.** Fold change in *ROSALIND* expression upon its inhibition with ASO#2 in MM383, MM047 and MM099. Significance was calculated by paired t-test. **C.** Western blot (left) and silver staining (right) of MM383, MM047 and MM099 mitoplast extracts upon inhibition of *ROSALIND*. Silver staining was used as a loading control. **D.** Densitometric analysis of DNP signal from the experiment in D. normalised on the silver staining. Paired t-test was used to calculate significance.

**Figure S6A-F:**
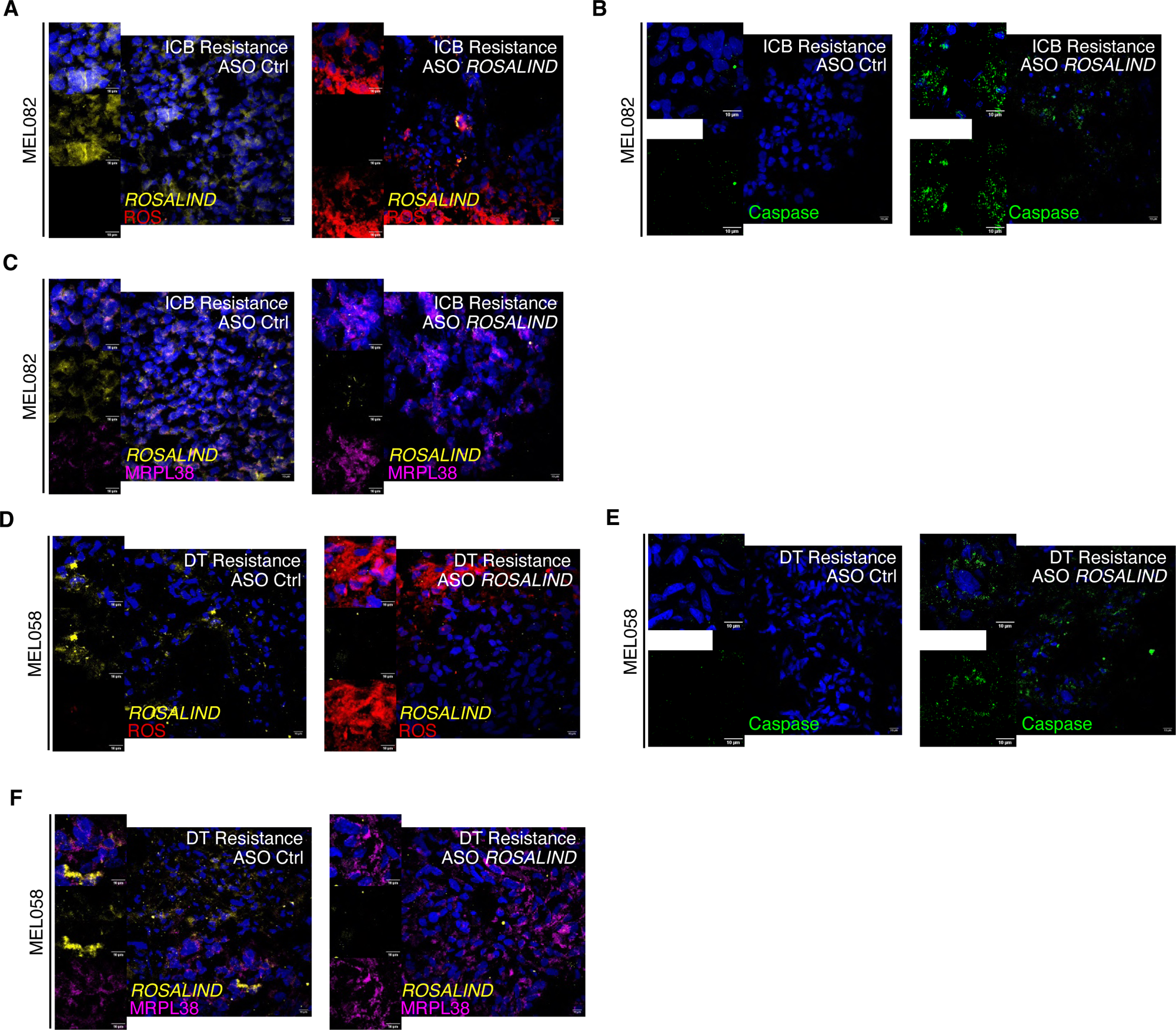
*ROSALIND* overcomes therapy resistance *ex vivo*. **A.** Representative confocal images of the staining for ROS (red) and for *ROSALIND (*by FISH, yellow) upon inhibition of *ROSALIND* in a PDXF model of MEL082 PDX derived from a patient resistant to immune therapy. **B.** Representative confocal images of the PDXF described in A. stained by immunofluorescence for caspase (green). **C.** Representative confocal images of the PDXF described in A. stained by FISH for *ROSALIND* (yellow) and immunofluorescence for MRPL38 (purple). **D.** Representative confocal images of the staining for ROS (red) and for *ROSALIND (*by FISH, yellow) upon inhibition of *ROSALIND* in a PDXF model of MEL058 PDX derived from a patient resistant to immune therapy. **E.** Representative confocal images of the PDXF described in G. stained by immunofluorescence for caspase (green). **F.** Representative confocal images of the PDXF described in G. stained by FISH for *ROSALIND* (yellow) and immunofluorescence for MRPL38 (purple).

